# Zebrafish Cre/*lox* regulated UFlip alleles generated by CRISPR/Cas targeted integration provide cell-type specific conditional gene inactivation

**DOI:** 10.1101/2021.06.18.448732

**Authors:** Maira P. Almeida, Sekhar Kambakam, Fang Liu, Zhitao Ming, Jordan M. Welker, Wesley A. Wierson, Laura E. Schultz-Rogers, Stephen C. Ekker, Karl J. Clark, Jeffrey J. Essner, Maura McGrail

## Abstract

The ability to regulate gene activity spatially and temporally is essential to investigate cell type specific gene function during development and in postembryonic processes and disease models. The Cre/*lox* system has been widely used for performing cell and tissue-specific conditional analysis of gene function in zebrafish, but simple and efficient methods for isolation of stable, Cre/*lox* regulated alleles are lacking. Here we applied our GeneWeld CRISPR/Cas9 short homology-directed targeted integration strategy to generate floxed conditional alleles that provide robust gene knockdown and strong loss of function phenotypes. A universal targeting vector, UFlip, with sites for cloning short 24-48 bp homology arms flanking a floxed mRFP gene trap plus secondary reporter cassette, was integrated into an intron in *hdac1, rbbp4*, and *rb1*. Active, gene off orientation *hdac1-UFlip-Off* and *rb1-UFlip-Off* integration alleles result in >99% reduction of gene expression in homozygotes and recapitulate known indel loss of function phenotypes. Passive, gene on orientation *rbbp4-UFlip-On* and *rb1-UFlip-On* integration alleles do not cause phenotypes in trans-heterozygous combination with an indel mutation. Cre recombinase injection leads to recombination at alternating pairs of *loxP* and *lox2272* sites, inverting and locking the cassette into the active, gene off orientation, and the expected mutant phenotypes. In combination with our endogenous neural progenitor Cre drivers we demonstrate *rbbp4-UFlip-On* and *rb1-UFlip-On* gene inactivation phenotypes can be restricted to specific neural cell populations. Replacement of the UFlip mRFP primary reporter gene trap with a 2A-RFP in *rbbp4-UFlip-Off*, or 2A-KalTA4 in *rb1-UFlip-Off*, shows strong RFP expression in wild type or UAS:RFP injected embryos, respectively. Together these results validate a simplified approach for efficient isolation of highly mutagenic Cre/*lox* responsive conditional gene alleles to advance zebrafish Cre recombinase genetics.

## Introduction

The ability to regulate gene activity spatially and temporally is essential to investigate cell type specific gene function during development and in postembryonic processes and disease models. Conditional gene inactivation can be achieved with the Cre/*lox* system in which the bacteriophage Cre recombinase promotes site-specific recombination at compatible *loxP* sites engineered in a gene of interest (Sauer and Henderson, 1988). Effective application of Cre/*lox* in vertebrates requires the isolation of transgenic lines expressing Cre, and gene modification to introduce *loxP* sequences at desired locations. Highly efficient homologous recombination driven site-directed genetic modification in embryonic stem cells enabled creation of the mouse Cre/*lox* resource that contains more than 2500 Cre drivers and *lox* conditional gene alleles (Bult et al., 2019). In zebrafish the lack of embryonic stem cell technology and inefficient homologous recombination in the early embryo necessitated alternative approaches to generate Cre/*lox* tools (Carney and Mosimann, 2018). Tol2 transposon transgenesis (Balciunas et al., 2006; Kawakami et al., 2000) combined with defined promoters (Mosimann et al., 2011) or BAC recombineering (Forster et al., 2017) has been used to develop Cre lines, and a rich source of zebrafish cell type specific Cre drivers has been generated by random Tol2 transposon enhancer trap screens (Jungke et al., 2015; Marquart et al., 2015; Tabor et al., 2019; Zhong et al., 2019). Cre/*lox* regulated conditional control of gene function in zebrafish was previously limited to generation of knock out alleles by random Tol2 transposon insertional mutagenesis with a floxed/FRT gene trap, which can be reverted by Cre or Flip mediated recombination (Clark et al., 2011; Ni et al., 2012; Trinh le et al., 2011). Only recently have gene editing methods been developed in zebrafish to simplify Cre driver isolation (Almeida et al., 2021; Kesavan et al., 2018) and generate genuine *lox*-regulated conditional alleles (Burg et al., 2018; Han et al., 2021; Hoshijima et al., 2016; Li et al., 2019; Sugimoto et al., 2017) by targeted integration. These advances in zebrafish Cre/*lox* genetics are being driven by rapidly evolving methods for homology directed gene editing using CRISPR/Cas9 and TALEN endonucleases.

We and others recently demonstrated CRISPR/Cas9 targeted integration is an efficient method to isolate zebrafish proneural specific Cre and CreERT2 drivers that are expressed under the control of endogenous gene regulatory elements (Almeida et al., 2021; Kesavan et al., 2018). Efforts to introduce *loxP* sequences directly into the zebrafish genome by somatic gene targeting was first described using TALENs to target a double strand break in the *cdrh2* gene, followed by repair from a DNA oligonucleotide template containing homologous sequences flanking a *loxP* site (Bedell et al., 2012). This approach was expanded upon using CRISPR/Cas9 to create a floxed allele in two2 zebrafish genes by sequential *loxP* oligonucleotide targeting at sites flanking an exon, and demonstrated robust gene knockdown and loss of function after Cre-mediated exon excision (Burg et al., 2018). Although this approach is effective for creating a Cre/*lox* regulated allele, a single integration event of a floxed cassette with a linked reporter has distinct advantages, including only one generation of targeting and fluorescent genotyping of the integration allele. The first example of a germline floxed conditional allele with linked reporter used a single TALEN site downstream of an exon to replace the endogenous exon by homologous recombination from a linear cassette containing 1 kb long homology arms flanking a floxed exon plus reporter (Hoshijima et al., 2016). A similar strategy using long homology arms was used to isolate a zebrafish conditional gene trap *shha* allele by integration of a modified *Flex* invertible gene trap (Ni et al., 2012), *Zwitch*, by homologous recombination from an intact plasmid (Sugimoto et al., 2017). Stable inversion of the *Zwitch* cassette by Cre led to effective *shha* gene knockdown and defective *shha* signaling. More recently homology independent targeting strategies have been described to generate conditional knockouts with a linked reporter, or bidirectional knockin of a dual reporter gene trap cassette, providing a method for labeling conditional gene knockout cells (Han et al., 2021; Li et al., 2019). As an alternative to isolation of stable germline conditional alleles, rapid tissue specific gene knockdown and cell labeling can be achieved by somatic targeting with a floxed Cas9-2A-GFP; U6:gRNA transposon in a Cre-specific transgenic background, allowing for simultaneous bi-allelic inactivation with GFP cell labeling (Hans et al., 2021). Together, these approaches provide a variety of effective conditional gene knockdown strategies with varying degrees of complexity in design and execution.

To address the need for a simple method to efficiently recover robust germline conditional alleles for any gene in zebrafish, here we applied our GeneWeld CRISPR/Cas9 targeted integration strategy directed by short homology (Wierson et al., 2020) to generate floxed conditional alleles by intron targeting of a highly mutagenic gene trap in three zebrafish genes. We showed previously our strategy significantly simplifies targeting vector assembly and enhances the efficiency of on target integration (Almeida et al., 2021; Wierson et al., 2020). The universal vector, UFlip, is derived from the pGTag and pPRISM targeted integration vectors, with sites for cloning short homology arms flanking a cargo of interest and universal gRNA sites for efficient CRISPR/Cas9 cutting *in vivo*. Homology arm cloning sites flank a floxed cassette that contains a mRFP or KalTA4 primary reporter based on the RP2 gene trap (Clark et al., 2011; Ichino et al., 2020), plus heart or lens BFP secondary reporter for transgenic identification. Using 24 or 48 bp short homology arms to drive integration at unique intron target gRNA sites, we recovered precise germline UFlip integration alleles in *hdac1, rb1* and *rbbp4* with frequencies of 4-10%. Integration alleles in the active, gene off orientation knock down gene expression to >99% of wild type levels and recapitulate indel mutant phenotypes, demonstrating highly effective loss of function by primary transcript splicing into the gene trap. Alleles with UFlip in the passive, gene on orientation are phenotypically normal. Exposure to Cre leads to efficient inversion and expected phenotypes, and cell type specific Cre drivers restrict conditional gene inactivation to the expected cell populations. Our results demonstrate UFlip CRISPR targeted integration is an effective approach to generate zebrafish Cre/*lox* conditional alleles for cell type specific gene inactivation.

## Results

### UFlip, a universal vector to generate stable Cre/*lox* regulated conditional gene alleles by CRISPR/Cas9 targeted integration

The <u>U</u>niversal <u>F</u>lip (UFlip) vector was designed to be used with the GeneWeld strategy for CRISPR/Cas9 targeted integration driven by short homology (Wierson et al., 2020). The vector contains a cassette with a floxed gene trap plus secondary marker flanked by cloning sites for homology arms (HA) complementary to the genomic CRISPR target site, and universal gRNA sites (UgRNA) for *in vivo* liberation of the cassette after injection into zebrafish embryos (Figure 1A). The gene trap was derived from the gene-breaking Tol2 transposon RP2 which contains a splice acceptor followed by mRFP and the ocean pout antifreeze gene transcriptional terminator, and has previously been shown to knockdown gene expression to >99% of wild type levels (Clark et al., 2011; Ichino et al., 2020). Like our recently published endogenous Cre driver lines isolated by targeted integration of a 2A-Cre plus secondary marker cassette (Almeida et al., 2021), the linked secondary marker drives tissue specific expression of blue fluorescent protein (BFP) from the *Xenopus* gamma crystallin 1 (*gcry1*) lens promoter or the zebrafish myosin light chain 7 (*myl7*) cardiac muscle promoter, simplifying identification of an integration allele. Alternating pairs of *loxP* and *lox2272* sites flanking the cassette are designed to drive two Cre recombination events that invert and lock the cassette in place, a design strategy described previously to generate floxed conditional alleles in mice (Robles-Oteiza et al., 2015; Schnutgen et al., 2003) and zebrafish (Sugimoto et al., 2017). *rox* sites flanking the entire cassette provide an alternative method for inversion using Dre recombinase (Figure 1A). Integration of UFlip into an intron in the active, gene off orientation is expected to lead to premature transcriptional termination of the primary transcript in the gene trap and splicing of upstream exons into mRFP (Figure 1B), resulting in a loss of function allele. In contrast, integration of UFlip into an intron in the passive, gene on orientation is not predicted to interrupt endogenous gene expression, since RNA polymerase will read through the cassette in the intron, which is then spliced out of the mature transcript (Figure 1C). For clarity, the UFlip alleles described here are referred to as *gene-UFlip-Off* when the cassette was integrated in the active, gene off orientation, and *gene-UFlip-On* when integrated in the passive, gene on orientation.

**Figure 1.**
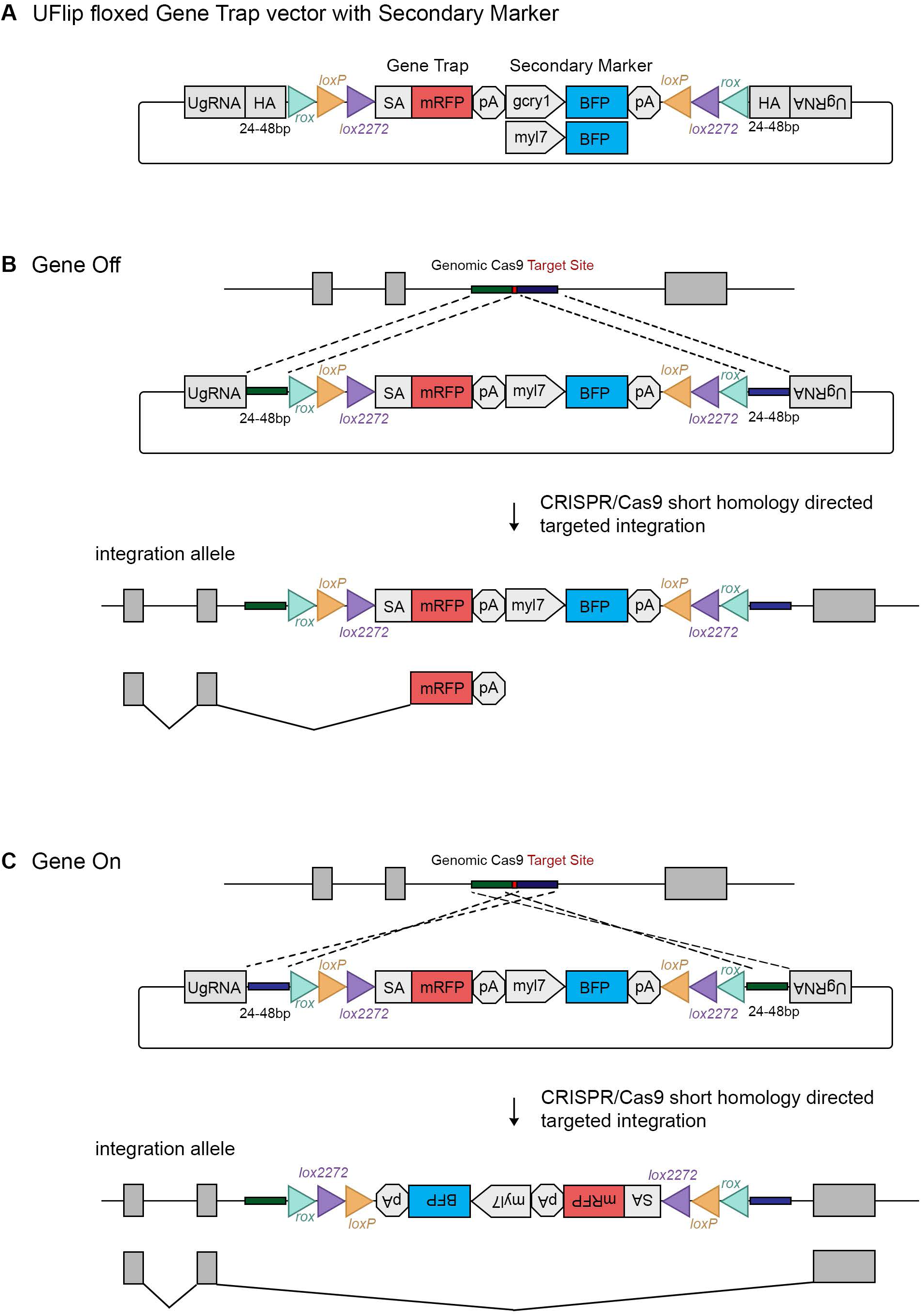
The UFlip floxed gene trap vector for isolation of conditional gene alleles generated by GeneWeld CRISPR/Cas9 targeted integration. (**A**) Diagram of the UFlip. The vector contains a floxed *rox loxP lox2272* gene trap plus secondary marker *loxP lox2272 rox* cassette. The cassette is flanked by cloning sites for homology arms (HA) complementary to a genomic CRISPR target site, and universal gRNA sites (UgRNA) for *in vivo* liberation of the targeting cassette. (**B**) Gene Off alleles are generated by integration of the UFlip cassette into an intron in the active orientation, leading to transcription termination and splicing of the primary transcript in the mRFP gene trap. (**C**) Gene On alleles are generated by integration of the UFlip cassette into an intron in the passive orientation. This is driven by cloning the genomic 5’ homology arm downstream of the UFlip cassette, and cloning the genomic 3’ homology arm upstream of the UFlip cassette. Integration at the genomic Cas9 target site occurs in the opposite orientation. During transcription RNA polymerase reads through the integrated UFlip cassette, which is then splice out with the intron during processing of the primary transcript. BFP, blue fluorescent protein; *gcry1*, gamma crystallin 1 promoter; *myl7*, cardiac myosin light chain 7 promoter; mRFP, monomeric red fluorescent protein; pA, transcription termination and polyadenylation signal; SA, splice acceptor.

### Isolation of *hdac1, rb1* and *rbbp4* UFlip conditional alleles

To test the functionality of zebrafish conditional alleles generated by UFlip targeted integration, we chose to target three genes, histone deacetylase 1 (*hdac1*), retinoblastoma 1 (*rb1*), and retinoblastoma binding protein 4 (*rbbp4*), for which we had previously isolated indel mutations that show strong loss of function phenotypes at the morphological and cellular level (Schultz et al., 2018; Solin et al., 2015). Intronic genomic DNA was amplified from wild type WIK fish, cloned, sequenced and analyzed to locate unique Cas9 gRNA sites that were shared among fish and that did not map to repetitive elements or retrotransposons. Cas9 target sites were identified in *hdac1* intron 5, *rb1* intron 6, and *rbbp4* intron 4 (Supplemental Table 1) and synthetic guides ordered from Synthego (https://www.synthego.com). Efficient indel formation at the intronic gRNA sites was confirmed by co-injection of gRNA and Cas9 mRNA into single cell embryos, followed by extraction of genomic DNA at 2 dpf or from adult fin clips. A PCR amplicon surrounding the target site was directly sequenced followed by ICE analysis (https://ice.synthego.com/#/). The *hdac1* intron 5, *rb1* intron 6 and *rbbp4* intron 4 guides showed 52%, 95% and 50% indel formation at the target site, respectively (Supplemental Figures 1, 2).

**Table 1.** Recovery of *hdac1, rb1*, and *rbbp4* UFlip floxed conditional alleles

| UFlip allele | Targeted Intron | Homology arm length<br>bp/bp | Injected F0 embryo secondary<br>marker expression | F0 germline transmission of<br>secondary marker | F0 germline transmission of<br>precise integration |
| --- | --- | --- | --- | --- | --- |
| *hdac1-i5-UFlip-Off* | 5 | 24/24 | 41% (30/73) | 30% (3/10) | 10% (1/10) |
| *hdac1-i5-UFlip-On* | 5 | 24/24 | 46% (85/185) | 12.5% (3/24) | 0% (0/24) |
| *rb1-i6-UFlip-Off* | 6 | 48/48 | 47% (34/73) | 9% (3/33) | 6% (2/33) |
| *rb1-i6-UFlip-On* | 6 | 48/48 | 43% (61/101) | 20% (5/25) | 4% (1/25) |
| *rbbp4-i4-UFlip-Off* | 4 | 48/48 | 79% (15/19) | 30% (3/10) | 0% (0/10) |
| *rbbp4-i4-UFlip-On* | 4 | 48/48 | 77% (23/30) | 36% (4/11) | 9% (1/11) |

**Figure 2.**
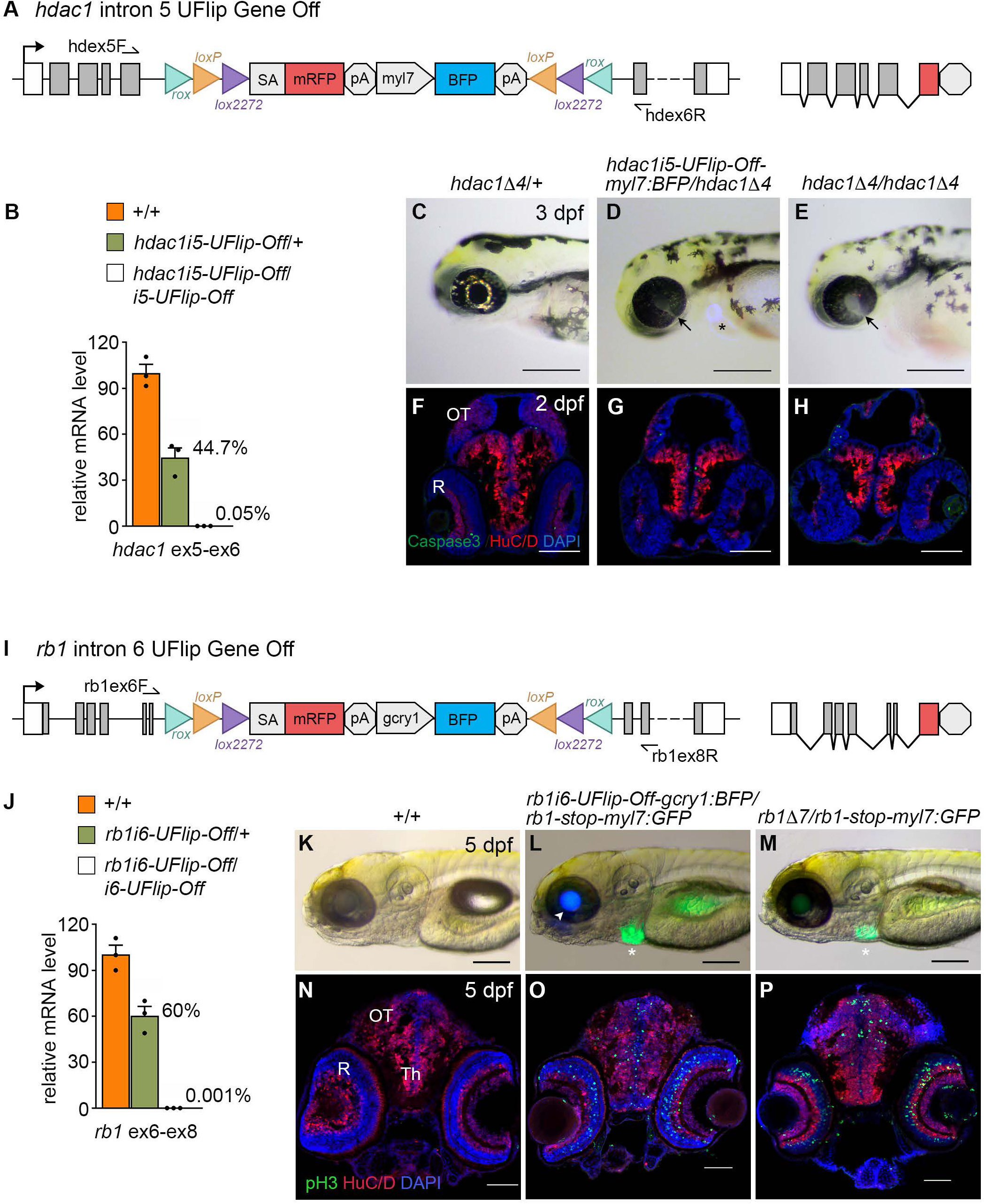
*hdac1-UFlip-Off* and *rb1-UFlip-Off* integration alleles result in >99% gene knockdown and recapitulate loss of function phenotypes. (**A**) Diagram of *hdac1-UFlip-Off* allele with location of exon 5 and exon 6 primers. (**B**) Quantitative RT-PCR of 3 dpf control +/+, *hdac1-UFlip-Off/+* and *hdac1-UFlip-Off/hdac1-UFlip-Off* larvae shows 99.95% knockdown of *hdac1* mRNA in homozygotes. (**C**-**H**) 3dpf gross phenotype and 2 dpf head tissue sectioned and labeled with Caspase-3a and HuC/D from *hdac1D4*/+ (**C, F**), *hdac1-UFlip-Off/hdac1D4* (**D, G**) and *hdac1D4*/*hda1cD4* larvae (**E, H**). Both *hdac1-UFlip-Off/hdac1D4* and *hdac1D4*/*hda1cD4* larvae show coloboma in the retina (**D, E** arrows) and reduced size of the midbrain (**G, H**). asterisk, BFP expression in the heart from the *hdac1-UFlip-Off* allele (**D**). (**I**) Diagram of *rb1-UFlip-Off* allele with location of exon 6 and exon 8 primers. (**J**) Quantitative RT-PCR of 5 dpf control +/+, *rb1-UFlip-Off/+* and *rb1-UFlip-Off/rb1-i6-UFlip-Off* larvae shows 99.99% knockdown of *rb1* mRNA in homozygotes. (**K**-**P**) 5 dpf gross phenotype and head tissue sectioned and labeled with pH3 and HuC/D from +/+ (**K, N**), *rb1-UFlip-Off/rb1-stop* (**L, O**) and *rb1D7*/*rb1-stop* larvae (**M, P**). Both *rb1-UFlip-Off/rb1-stop* and *rb1D7*/*rb1-stop* larvae do not develop swim bladders (**L, M**) and show proliferating pH3 positive cells throughout the midbrain optic tectum and retina (**O, P**). Arrowhead points to lens BFP expression from the *rb1-UFlip-Off* allele, asterisks mark heart GFP expression from the *rb1-stop* allele (**L, M**). Th, thalamic region; OT, optic tectum; R, retina. Scale bars: 250 μm (K-M); 200 μm (C-E); 100 μm (F-H); 50 μm (**N**-**P**).

UFlip targeting vectors were assembled with 5’ and 3’ homology arms complementary to the DNA flanking the Cas9 genomic DNA double strand break site. The *hdac1-UFlip-Off* and *-On* vectors contained 24 bp 5’ and 3’ homology arms, while the *rb1-UFlip-Off* and *-On* vectors, and the *rbbp4-UFlip-Off* and *-O*n vectors, contained 48 bp 5’ and 3’ homology arms (Supplemental Table 1). One cell stage WIK embryos were co-injected with targeting vector (10 pg), Cas9 mRNA (150 pg), universal gRNA (25 pg), and gene specific gRNA (25 pg). Four to six embryos with positive expression of the secondary lens or heart reporter were selected to test for evidence of on target integration by PCR amplification of the 5’ and 3’ junctions (data not shown), and secondary reporter positive siblings were raised to adulthood. Germline transmission of integration alleles with positive secondary marker expression ranged from 9% (3/33) to 36% (4/11). Fin clips from F1 adults were sequenced to determine the rate of recovery of precise targeted integration alleles (Supplemental Figure 3). The final transmission rates of alleles with both 5’ and 3’ precise junctions were *hdac1-UFlip-Off* 10% (1/10), *rb1-UFlip-Off* 6% (2/33), *rb1-UFlip-On* 4% (1/25), and *rbbp4-UFlip-On* 9% (1/11) (Table 1). 1/25 founders transmitted an *hdac1-UFlip-On* integration, but neither 5’ nor 3’ junctions were amplified by PCR. Injection of Dre mRNA into *hdac1-UFlip-Off*/+ embryos led to robust inversion of the cassette by Dre/*rox* mediated recombination (Supplemental Figure 4), providing a possible alternative approach for recovery of a *hdac1-UFlip-On* allele in the next generation. F1 adults with precise integrations were outcrossed to WIK to establish F2 families of the alleles *hdac1-UFlip-Off*^*is71*^, *rb1-UFlip-Off*^*is58*^, *rb1-UFlip-On*^*is57*^, and *rbbp4-UFlip-On*^*is61*^.

**Figure 3.**
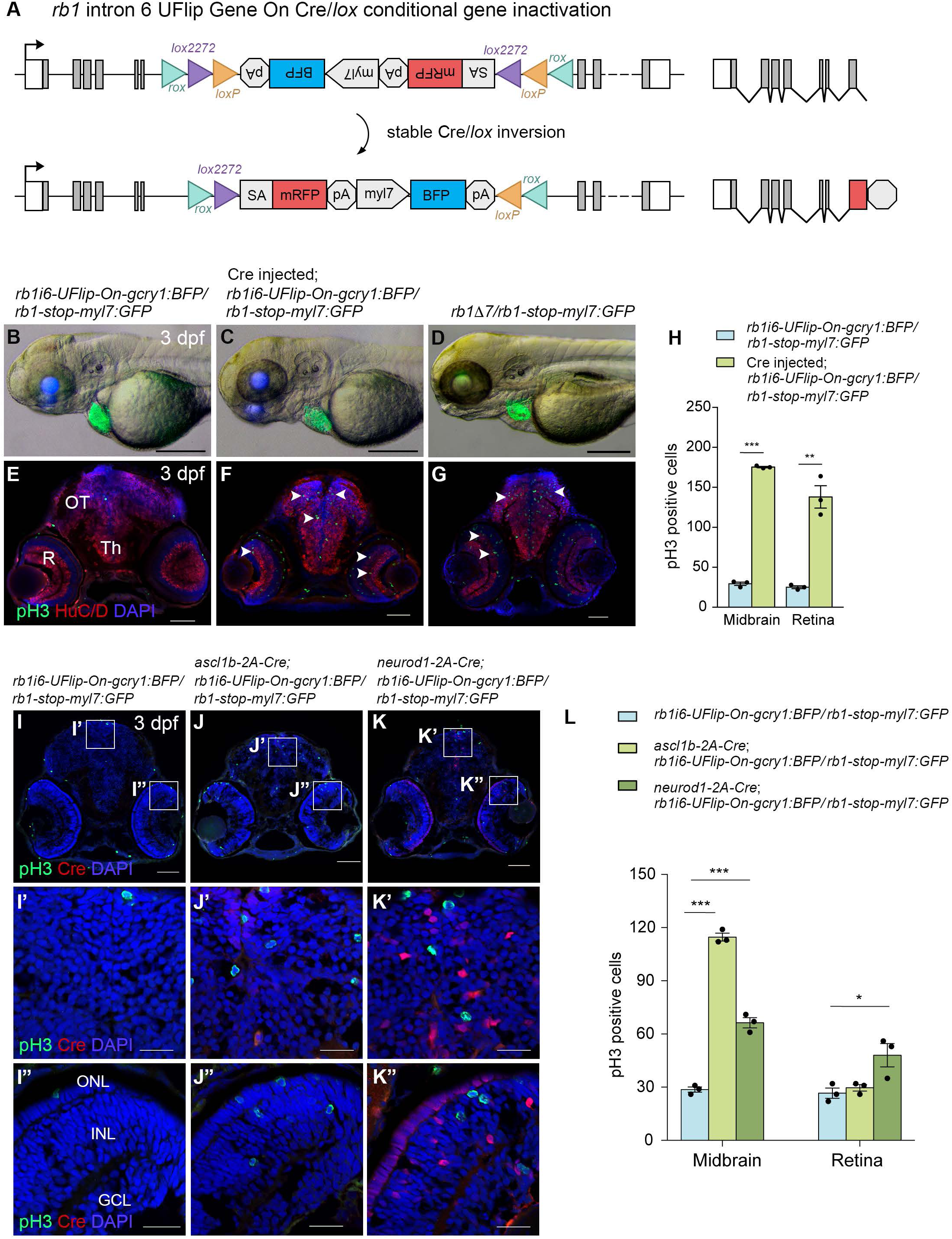
Cre-mediated recombination of the *rb1-UFlip-On* allele leads to conditional gene inactivation. (**A**) Diagram of *rb1-UFlip-On* allele before and after stable Cre/*lox* inversion. (**B**-**D**) 3 dpf gross phenotype of control uninjected *rb1-UFlip-On/rb1-stop* (**B**), Cre injected *rb1-UFlip-On/rb1-stop* (**C**), and uninjected *rb1D7 /rb1-stop* (**D**) larvae. pH3 and HuC/D immunolabeling of 3 dpf head tissue from control uninjected *rb1-UFlip-On/rb1-stop* (**E**), Cre injected *rb1-UFlip-On/rb1-stop* (**F**), and uninjected *rb1D7/rb1-stop* (**G**) larvae. (**H**) Quantification bar graph of pH3 positive cells in *rb1-UFlip-On/rb1-stop* and Cre injected *rb1-UFlip-On/rb1-stop* larvae (n=3 biological replicates, with 3 sections counted for each replicate) in midbrain (*** *p*<0.0001) and retina (** *p*<0.001). (**I**-**K**) 3 dpf *rb1-UFlip-On/rb1-stop* (**I**), *ascl1b-2A-Cre; rb1-UFlip-On/rb1-stop* (**J**), and *neurod1-2A-Cre; rb1-UFlip-On/rb1-stop* (**K**) labeled with pH3 and Cre antibodies. Rectangles indicate higher magnification images of midbrain (**I**’, **J**’, **K**’) and retina (**I**”, **J**”, **K**”). (**L**) Quantification bar graph of pH3 positive cells in midbrain and retina of *rb1-UFlip-On/rb1-stop, ascl1b-2A-Cre; rb1-UFlip-On/rb1-stop* (midbrain *** *p*<0.0001), and *neurod1-2A-Cre; rb1-UFlip-On/rb1-stop* (midbrain *** *p*<0.0001; retina * *p*<0.04) larvae (n=3 biological replicates, with 3 sections counted for each replicate). Graph shows mean ± s.e.m. (two tailed unpaired Student’s t-test). GCL, ganglion cell layer; INL, inner nuclear layer; ONL, outer nuclear layer; OT, midbrain optic tectum; R, retina; Th, midbrain thalamic region. Scale bars: 200 μm (**B**-**D**); 50 μm (**E**-**G, I-K**); 20 μm (**I’**-**K”**).

**Figure 4.**
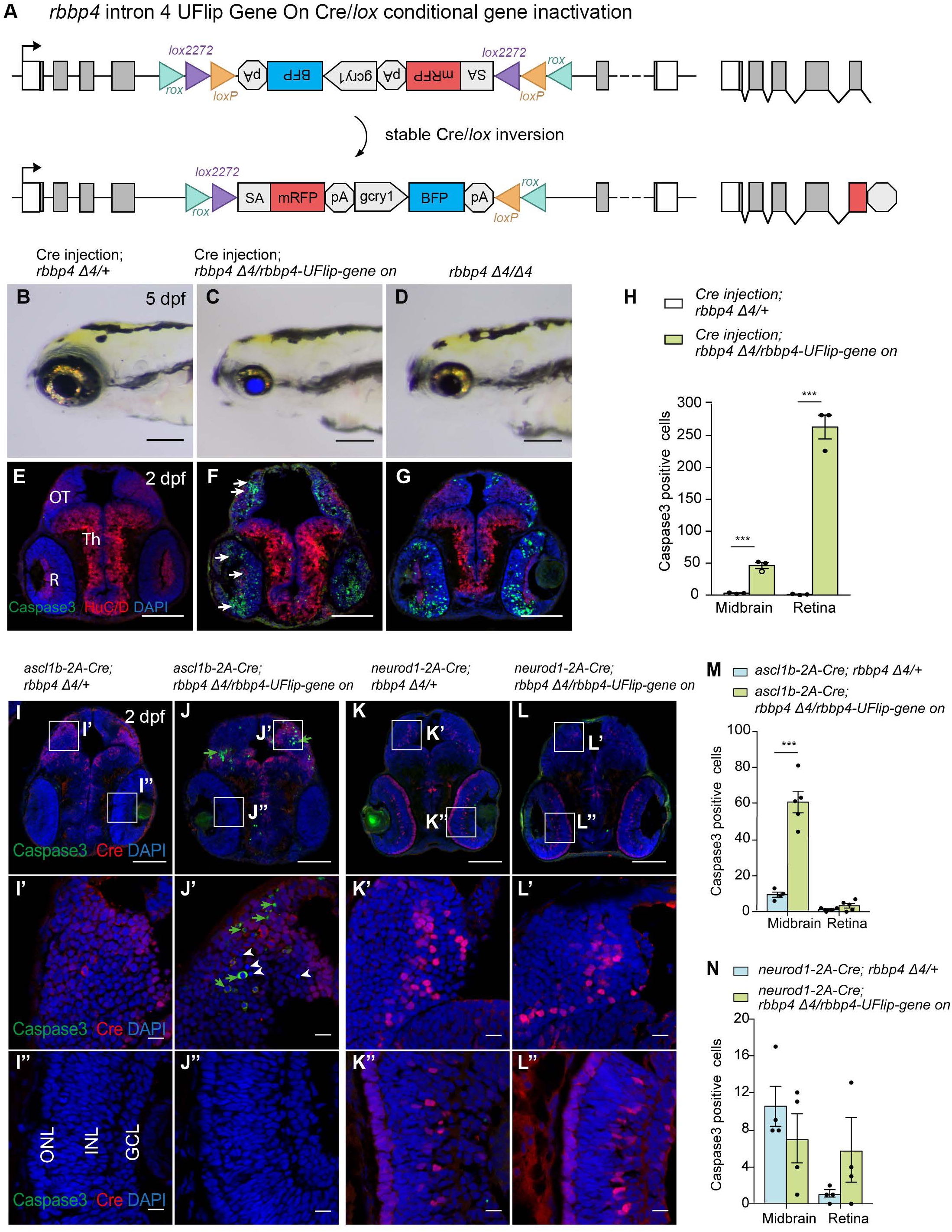
Cre-mediated recombination of the *rbbp4-UFlip-On* allele leads to conditional gene inactivation. (**A**) Diagram of *rbbp4-UFlip-On* allele before and after stable Cre/*lox* inversion. (**B**-**D**) 5 dpf gross phenotype of Cre injected control *rbbp4D4/+* (**B**), Cre injected *rbbp4D4/rbbp4-UFlip-On* (**C**), and uninjected *rbbp4D4/rbbp4D4* (**D**) larvae. Caspase-3a and HuC/D immunolabeling of 2 dpf head tissue from control uninjected *rbbp4D4/+* (**E**), Cre injected *rbbp4D4/rbbp4-UFlip-On* (**F**), and uninjected *rbbp4D4/rbbp4D4* (**G**) embryos. Arrows point to Caspase-3a labelled cells (**F**). (**H**) Quantification bar graph of Caspase-3a positive cells in Cre injected control *rbbp4D4/+* and *rbbp4D4/rbbp4-UFlip-On* embryos (n=3 biological replicates, with 3 sections counted for each replicate) in midbrain (*** *p*<0.0008) and retina (*** *p*<0.0002). (**I**-**L**) 2 dpf *ascl1b-2A-Cre; rbbp4D4/+* (**I**), *ascl1b-2A-Cre; rbbp4D4/rbbp4-UFlip-On* (**J**), *neurod1-2A-Cre; rbbp4D4/+* (**K**), and *neurod1-2A-Cre; rbbp4D4/rbbp4-UFlip-On* (**L**) sections labeled with Caspase-3a and Cre antibodies. Rectangles indicate higher magnification images of midbrain (**I**’, **J**’, **K**’, **L**) and retina (**I**”, **J**”, **K**”, **L”**). Green arrows indicate Caspase-3a positive cells; white arrowheads mark pyknotic nuclei (**J’**). (**M**) Quantification bar graph of Caspase-3a positive cells in *ascl1b-2A-Cre; rbbp4D4/+* and *ascl1b-2A-Cre; rbbp4D4/rbbp4-UFlip-On* embryos (n=3 biological replicates, with 3 sections counted for each replicate) in midbrain (*** *p*<0.0002) and retina. (**N**) Quantification bar graph of Caspase-3a positive cells in *neurod1-2A-Cre; rbbp4D4/+* and *neurod1-2A-Cre; rbbp4D4/rbbp4-UFlip-On* embryos (n=3 biological replicates, with 3 sections counted for each replicate) in midbrain and retina. Graph shows mean ± s.e.m. (two tailed unpaired Student’s t-test). GCL, ganglion cell layer; INL, inner nuclear layer; ONL, outer nuclear layer; OT, midbrain optic tectum; R, retina; Th, midbrain thalamic region. Scale bars: 200 μm (**B**-**D**); 100 μm (**E**-**G, I**-**L**); 10 μm (**I’**-**L”**).

### *hdac1-UFlip-Off* and *rb1-UFlip-Off* alleles provide robust gene knockdown and loss of function

To demonstrate that integration of the UFlip cassette leads to effective gene knockdown we examined gene expression by quantitative PCR in homozygotes and tested for mutant phenotypes in trans-heterozygotes with established loss of function alleles. Homozygous *hdac1-UFlip-Off* 3dpf larvae show a >99% reduction in *hdac1* mRNA levels in comparison to wild type sibling larvae (Figure 2 A, B). In comparison to wild type 3 dpf larvae, *hdac1-UFlip-Off*/*hdac1Δ4* trans-heterozygotes have a smaller head and retinal coloboma (Figure 2 C, D arrow), similar to *hdac1Δ4/hdac1Δ4* homozygotes (Figure 2, E arrow) (Schultz et al., 2018). The *hdac1-UFlip-Off* secondary reporter *myl7:BFP* expression is visible in the heart (Figure 2 D asterisk). 2 dpf sectioned head tissue labeled with antibodies to activated Caspase-3a and the neuronal marker HuC/D revealed that in comparison to wild type, *hdac1-UFlip-Off*/*hdac1Δ4* show a reduced optic tectum, disorganized retina, but minimal apoptosis (Figure 2 F, G), like the *hdac1Δ4/hdac1Δ4* homozygous phenotype (Figure H).

Quantitative RT-PCR of homozygous *rb1-UFlip-Off* 3 dpf larvae also showed >99% knockdown of *rb1* mRNA levels in comparison to wild type sibling larvae (Figure 2 I, J). To facilitate identification of trans heterozygous *rb1* mutant larvae, we isolated an *rb1* exon 2 CRISPR integration allele, *rb1-stop-myl7:GFP*^*is59*^ (*rb1-stop*) using a targeting vector design similar to our recently published method to isolate endogenous Cre lines in proneural genes (Almeida et al., 2021). In the *stop-myl:GFP* cassette the Cre cDNA was replaced with a sequence containing three translation termination codons, in each of the three reading frames, followed by a transcription terminator plus a *myl7:GFP* secondary marker (Supplemental Figure 5 A, B). The *rb1-stop* allele is larval lethal in combination with *rb1Δ7* and shows excess proliferation throughout the midbrain and retina (Supplemental Figure 5 C), similar to our previous observations of *rb1Δ7*/*rb1Δ7* homozygotes (Schultz et al., 2018). *rb1-UFlip-Off/rb1-stop* 5 dpf larvae appear morphologically normal compared to wild type with the exception of the lack of a swim bladder (Figure 2 K-L), like *rb1Δ7*/*rb1-stop* larvae (Figure 2 M). Both secondary reporters from the *rb1-UFlip-Off* allele (lens BFP; Figure 2 L arrowhead) and the *rb1-stop* allele (heart GFP; Figure 2 L asterisk) are visible, illustrating the utility of integrating linked reporters for allele identification. Compared to wild type, sectioned head tissue from 5 dpf larvae shows excess proliferation in the midbrain and retina in *rb1-UFlip-Off/rb1-stop*, similar to *rb1Δ7/rb1-stop* trans-heterozygotes (Figure 2 N-P). Together these results demonstrate the active UFlip integration alleles provide robust gene knockdown and recapitulate loss of function phenotypes.

### *UFlip-2A-mRFP-Off* and *-2A-KalTA4-Off* integration allows detection of the gene trap fluorescent primary reporter

The results above demonstrate integration of the UFlip allele in the gene off orientation leads to robust gene knockdown and indicate the splice acceptor and primary reporter mRFP gene trap are functioning as expected to cause premature transcription termination. We confirmed splicing of the primary transcript into the gene trap using RT-PCR for both *hdac1-UFlip-Off* and *rb1-UFlip-Off* alleles (data not shown), however neither showed mRFP reporter expression in embryos or larvae, possibly due to upstream endogenous polypeptide sequences disrupting mRNA activity. To address this, the mRFP cDNA in the UFlip cassette was replaced with 2A-mRFP or 2A-KalTA4 cDNAs preceded by the porcine teschovirus-1 polyprotein 2A peptide to release the reporter from upstream sequences. *hdac1-UFlip-2A-KalTA4-Off* and *rb1-UFlip-2AKalTA4-Off* constructs were injected into the UAS reporter line *Tg*(*Tol2*<*14XUAS:mRFP; gcry1:GFP*>) (Balciuniene et al., 2013), and both led to broad mRFP expression in injected larvae (Supplemental Figures 6, 7). 20% (24/121) of *hdac1-UFlip-2A-KalTA4-Off* injected embryos showed mRFP throughout the nervous system (Supplemental Figure 6 A-C). PCR amplification and sequencing of genomic DNA-UFlip junctions revealed on target integration in mRFP positive embryos (Supplemental Figure 6 D). Similarly, 41% (22/54) of *rb1-UFlip-2AKalTA4-Off* injected embryos showed widespread mRFP throughout larvae and evidence of on target integration by PCR analysis of genomic DNA-UFlip junctions (Supplemental Figure 7 A-D). Injection of *rbbp4-UFlip-2A-mRFP-Off* into wild type WIK resulted in mRFP expression throughout the nervous system in 16-20% of injected embryos and confirmation of on target integration by PCR and sequence analysis (Supplemental Figure 8 A-D). Together the data indicates detection of the primary reporter is impacted by intronic target site and the level of expression of the endogenous gene.

### Cre/*lox* recombination of *rb1-UFlip-On* allele leads to efficient conditional gene inactivation and recapitulates morphological and cellular loss of function phenotypes

To demonstrate the ability of UFlip gene on integration alleles to create conditional gene inactivation in response to Cre-mediated recombination we first used Cre mRNA injection into mutant embryos. 150 pg Cre mRNA was injected into embryos from a cross between heterozygous *rb1-UFlip-On/+* and *rb1-stop/+* adults to induce UFlip cassette inversion from the passive to the active orientation (Figure 3 A). At 3 dpf *rb1-UFlip-On*/*rb1-stop* uninjected and injected larvae appear morphologically normal (Figure 3 B, C) like *rb1Δ7*/*rb1-stop* trans-heterozygotes (Figure 3 D). However, in comparison to uninjected control sectioned head tissue (Figure 3 E), Cre injected *rb1-UFlip-On*/*rb1-stop* showed phosphohistone pH3 labeling throughout the brain and retina (Figure 3 F arrowheads), similar to the loss of function phenotype of *rb1Δ7*/*rb1-stop* (Figure 3 G). Quantification showed a significant increase in the number of pH3 positive cells in injected larvae in both the midbrain (*p*<0.0001) and retina (*p*<0.001) (Figure 3 H). Stable inversion of the UFlip cassette in Cre injected *rb1-UFlip-On*/*rb1-stop* embryos was confirmed by PCR junction analysis and sequencing of the junction amplicons (Supplemental Figure 9 A-F). These results demonstrate Cre recombination leads to stable inversion of the UFlip cassette and conditional gene inactivation which replicates the *rb1* loss of function phenotype at the cellular level.

### Cell type specific *rb1-UFlip-On* conditional gene inactivation reveals *rb1* requirement to suppress cell cycle entry in *ascl1b* and *neurod1* neural progenitors

To demonstrate the UFlip cassette can lead to cell type specific conditional gene inactivation in combination with Cre drivers that define distinct cell populations, we crossed *rb1-UFlip-On/+* with *rb1-stop/+* ; *ascl1b-2A-Cre/+* or *rb1-stop/+* ; *neurod1-2A-Cre/+* adults. In larva the neural progenitor specific *ascl1b-2A-Cre* allele expresses Cre specifically in the optic tectum but not in the retina, whereas *neurod1-2A-Cre/+* expresses Cre in committed progenitors/newborn neurons throughout the midbrain and retina (Almeida et al., 2021). 3 dpf control *rb1-UFlip-On*/*rb1-stop* (Figure 3 I-I”) and *ascl1b-2A-Cre/+; rb1-UFlip-On*/*rb1-stop* (Figure 3 J-J”) larvae were sectioned and labeled with antibodies to Cre and phosphohistone pH3 to correlate Cre expression with cell cycle entry. In *ascl1b-2A-Cre/+; rb1-UFlip-On*/*rb1-stop* embryos Cre was detected in cells in the dorsal region of the optic tectum and pH3 positive cells were detected in adjacent regions (Figure 3 J, J’). Neither Cre nor pH3 positive cells were detected in the retina (Figure 3 J, J”). In *neurod1-2A-Cre/+; rb1-UFlip-On*/*rb1-stop* embryos, Cre expression was visible in the midbrain and in the retinal ganglion cell, inner and outer nuclear layers of the retina. However, an increase in pH3 positive cells appeared to be restricted to the optic tectum (Figure 3 K, K’) not in the retina (Figure 3 K, K”). Quantification of pH3 positive cells confirmed a significant increase in the optic tectum (p<0.0001) but not in the retina after inactivation with *ascl1b-2A-Cre* (Figure 3 L). A similar significant increase in the number of pH3 positive cells after *neurod1-2A-Cre* inactivation was observed in the midbrain (*p*<0.0001), with a less significant increase in the retina (*p*<0.04) (Figure 3 L). Together these results indicate *rb1* is required to prevent cell cycle entry in *ascl1b* and *neurod1* neural progenitor populations in the developing midbrain. Overall, these results demonstrate Cre drivers can restrict UFlip gene inactivation and reveal gene requirements in specific cell populations.

### Cre/*lox* recombination of *rbbp4-UFlip-On* allele leads to efficient conditional gene inactivation and recapitulates morphological and cellular loss of function phenotypes

We validated cell type specific Cre-mediated UFlip conditional gene inactivation at a second locus using the *rbbp4-UFlip-On* allele (Figure 4 A). Cre mRNA was injected into embryos from a cross between *rbbp4-UFlip-On/+* and adults heterozygous for the *rbbp4* indel allele *rbbp4Δ4*^*is60*^ (Schultz et al., 2018). At 5 dpf control heterozygous *rbbp4Δ4*^*is60*^*/+* larvae do not show any morphological defects (Figure 4 B) whereas sibling *rbbp4Δ4*^*is60*^*/rbbp4-UFlip-On* trans-heterozygous larvae show microcephaly and microphthalmia (Figure 4 C), like the *rbbp4Δ4*^*is60*^*/rbbp4Δ4*^*is60*^ indel loss of function phenotype (Figure 4 D). Confirmation of stable inversion of the UFlip cassette and correlation of the *rbbp4Δ4*^*is60*^*/rbbp4-UFlip-On* genotype with phenotype were determined by PCR junction analysis and genotyping of the *rbbp4* locus (Supplemental Figure 10). Uninjected and injected larvae were sorted based on phenotype and expression of the UFlip allele secondary marker *gcry1:BFP*. 5’ and 3’ junctions of the inverted cassette were present only from injected embryos carrying the *rbbp4-UFlip-On* allele (Supplemental Figure 10 B, C, D). The gross *rbbp4* loss of function phenotype was only present in Cre mRNA injected *rbbp4-UFlip-On* larvae that also inherited the *rbbp4Δ4*^*is60*^ allele (Figure 9 B, E). Recapitulation of the *rbbp4* loss of function phenotype was also observed at the cellular level. Sectioned head tissue from Cre mRNA injected control and *rbbp4Δ4*^*is60*^*/rbbp4-UFlip-On* siblings was labeled with antibodies to activated Caspase-3a and revealed extensive apoptosis throughout the midbrain optic tectum and retinas (Figure 4 E, F arrows), similar to what was previously observed in *rbbp4Δ4*^*is60*^*/rbbp4Δ4*^*is60*^ homozygotes (Figure 4 G). Quantification of Caspase-3a labeling showed that *rbbp4* inactivation led to a significant increase in apoptosis in the midbrain (*p*<0.0008) and retina (*p*<0.0002) (Figure 4 H). These results demonstrate robust *rbbp4-UFlip-On* conditional inactivation replicates the *rbbp4* loss of function phenotype and shows its requirement for cell survival during neurogenesis.

### Cell type specific *rbbp4-UFlip-On* conditional gene inactivation reveals *rbbp4* requirement to suppress apoptosis in *ascl1b* but not *neurod1* neural progenitors

To test whether *rbbp4* is required to prevent apoptosis specifically in neural progenitors, *rbbp4-UFlip-On/+* fish were crossed with *rbbp4Δ4*^*is60*^*/+* ; *ascl1b-2A-Cre/+* adults to drive Cre recombination in *ascl1b* neural progenitors in the optic tectum. Sectioned head tissue from 2 dpf control *ascl1b-2A-Cre/+*; *rbbp4Δ4*^*is60*^*/+* and sibling *ascl1b-2A-Cre/+* ; *rbbp4Δ4*^*is60*^*/rbbp4-UFlip-On* embryos was labeled with antibodies to Cre and activated Caspase-3a. Cre expression was restricted to the dorsal region of the midbrain, and as expected, in contrast to control embryos which showed very little apoptosis (Figure 4 I-I”), *ascl1b-2A-Cre/+*; *rbbp4Δ4*^*is60*^*/rbbp4-UFlip-On* embryos showed Caspase-3a positive cells and disintegrating nuclei in the optic tectum (Figure 4 J, J’ arrows and arrowheads). Apoptosis was not detected in the retina (Figure 4 J, J”) as expected. *rbbp4* was inactivated in the *neurod1* cell population with the *neurod1-2A-Cre* Cre driver. Immunolabeling of control *neurod1-2A-Cre/+* ; *rbbp4Δ4is60/+* and sibling *neurod1-2A-Cre/+* ; *rbbp4Δ4*^*is60*^*/rbbp4-UFlip-On* sectioned head tissue showed Cre expression in the midbrain and in the retinal ganglion cell, inner and outer nuclear layers of the retina. Neither control embryos or *neurod1-2A-Cre/+* ; *rbbp4Δ4*^*is60*^*/rbbp4-UFlip-On* embryos showed extensive Caspase-3a labeling (Figure 4 K, L). Quantification of Caspase-3a positive cells revealed the level of apoptosis after *ascl1b-2A-Cre* mediated *rbbp4* inactivation significantly increased in the optic tectum (*p*<0.0002) but not in the retina (Figure L). In contrast, inactivation in the *neurod1-2A-Cre* population did not lead to a significant increase in apoptosis in either the midbrain or the retina (Figure M). In the *ascl1b* specific neural progenitor population, *rbbp4* is required for preventing apoptosis. In contrast, inactivation of *rbbp4* in the *neurod1* committed progenitor population did not lead to programmed cell death. Together the data show that *UFlip-On* alleles, in combination with defined Cre drivers, can restrict conditional gene inactivation to specific cell populations and reveal the requirement for gene function in distinct cell types.

## Discussion

In this study we used GeneWeld CRISPR/Cas9 targeting to isolate Cre/*lox* regulated conditional alleles with linked reporters by integration of a floxed gene trap cassette, UFlip, at a single gRNA location in an intron. Our results showed UFlip alleles led to >99% gene knockdown at the molecular level, robust Cre/*lox* mediated inversion and conditional inactivation, and recapitulated expected loss of function phenotypes at the morphological and cellular levels. We further demonstrated using neural progenitor Cre drivers that *rb1* and *rbbp4* conditional gene inactivation can be restricted to distinct neural cell populations, revealing cell type specific requirements for gene activity that are only inferred by loss of function homozygous indel mutation analysis. Our approach is simple in design, efficient, generates strong loss-of-function Cre/*lox* regulated alleles, and can be applied to any gene of interest.

We assembled UFlip vectors with 24 or 48 bp homology arms to generate both active (Gene Off) and passive (Gene On) integrations in an intron in three genes, *hdac1, rb1* and *rbbp4*. Of the 6 potential UFlip alleles, we recovered 4 precise on target integration alleles with a frequency of 4-10% of founders screened. This frequency is lower than our previous reports for GeneWeld germline integration alleles to tag genes with in frame reporters (22-100%) (Wierson et al., 2020) or generate endogenous 2A-Cre drivers (10-100%) (Almeida et al., 2021; Wierson et al., 2020). In those studies embryos showing expression of the fluorescent tag or Cre-mediated switch of a floxed reporter were selected and raised to adulthood to screen for germline transmission. In this report we selected embryos based on expression of the secondary reporter, which does not reflect on target integration. The lower frequency reported here may reflect that in the absence of a primary fluorescent reporter that allows for selection of injected embryos with on target integration, a greater number of adults must be screened to identify a founder transmitting an allele. Similar to our findings when isolating endogenous Cre drivers (Almeida et al., 2021), the UFlip allele founders transmitted both precise and imprecise integration events. We recovered *rb1-UFlip-On* and *rbbp4-UFlip-On* precise integration alleles at a frequency of 4% (1/25) and 9% (1/11), and *rb1-UFlip-Off* and *hdac1-UFlip-Off* alleles at a frequency of 6% (2/33) and 10% (1/10), a reasonable number of founders to screen. Previously reported methods for recovery of intron targeted floxed conditional alleles varied widely in frequency, from 12% for homology directed integration from a linear template with long 1-1.5 kb homology arms (Hoshijima et al., 2016), 53% by homologous recombination from a circular plasmid with 1 kb long homology arms (Sugimoto et al., 2017), to 56% – 60% for integration by Non-Homologous End Joining after preselection of a primary reporter (Han et al., 2021; Li et al., 2019). Our results demonstrate efficient recovery of precision integration alleles without requiring preselection of embryos expressing a fluorescent primary reporter, expanding the possibilities for novel integration cassette design and gene modification. The simplicity of UFlip vector construction and ability to recover precise integrations, in either orientation, underscore the power of our approach for isolation of stable Cre/*lox* regulated conditional alleles.

For effective conditional gene inactivation it is essential that the method leads to robust knockdown of wildtype mRNA expression and recapitulates loss of function phenotypes for genetic analysis. Our UFlip conditional construct is based on the Tol2 transposon RP2 gene trap, which contains a transcriptional terminator that effectively blocks transcription readthrough and reduces transcript levels by >99% (Clark et al., 2011; Ichino et al., 2020). The efficacy of gene knockdown by the UFlip gene trap was demonstrated by quantitative RT-qPCR and revealed >99% reduction in mRNA levels. Cre/*lox* mediated inversion of the UFlip cassette from the passive to active orientation was highly efficient after Cre mRNA injection or exposure to a transgenic source of Cre, generating expected loss of function phenotypes in combination with known indel mutations. While the recently published Cre-regulated CRISPR indel targeting study provides a rapid method for conditional mutagenesis and phenotypic knockout in embryos (Hans et al., 2021), this approach will be limited to genes that are not subject to genetic compensation. In contrast, a significant advantage of our stable conditional UFlip alleles rests on the strength of transcriptional termination by the UFlip cassette in the active orientation, which should eliminate activation of genetic compensation caused by frameshift mRNA non-sense mediated decay (El-Brolosy et al., 2019). We further validated effective UFlip conditional gene inactivation at the cellular level. We previously demonstrated that *rb1* is required to prevent neural progenitor cell cycle re-entry in the developing zebrafish brain and *rbbp4* is necessary for neural progenitor survival and differentiation (Schultz et al., 2018). Using our *ascl1b-2A-Cre* and *neurod1-2A-Cre* drivers in combination with *rb1-UFlip-On* and *rbbp4-UFlip-On* conditional alleles we demonstrated the requirement for each gene could be examined specifically in the *ascl1b* or *neurod1* neural progenitor populations. These genetic tools provide a foundation for future mechanistic studies investigating the interaction of *rb1* and *rbbp4* specifically in neural progenitor cell cycle regulation and differentiation, but can also be used with other cell type specific Cre drivers. Our results illustrate the robustness of the UFlip conditional alleles for cell type specific gene knockout analysis and their potential application in different developmental or disease model contexts.

To isolate a UFlip integration allele it was necessary to sequence the introns in our laboratory wild type WIK strain before gRNA design, to identify a unique gRNA site that was shared among adults in the WIK population and did not map to a retrotransposon or repetitive element. The *hdac1, rb1* and *rbbp4* indel or loss of function alleles we used to validate the UFlip conditional alleles were located in an early coding exon in the gene, except for *hdac1Δ4*. The first gRNAs meeting these criteria mapped in introns located downstream of the indel or loss of function alleles *hdac1Δ4* (exon 5), *rb1Δ7* or *rb1-stop* (exon 2), and *rbbp4Δ4* (exon 2). This may have accounted for the absence of expression of the gene trap mRFP primary reporter in the *hdac1-UFlip-Off* (intron 5) and *rb1-UFlip-Off* (intron 6) alleles, due to upstream Hdac1 and Rb1 polypeptide sequences interfering with in frame mRFP protein folding or expression. We updated the UFlip construct by replacing the mRFP in the gene trap with 2A-RFP or 2A-KalTA4, which resulted in robust primary reporter expression after targeting in Tol2<14XUAS:RFP> embryos injected with *hdac1-2AKalTA4-UFlip-Off* or *rb1-2AKalTA4-UFlip-Off*, or WIK embryos injected with *rbbp4-2AmRFP-UFlip-Off*. Integration alleles generated with the updated UFlip construct will allow fluorescent labeling of cells with Cre/*lox* conditional gene inactivation, as described in other recent studies (Han et al., 2021; Hans et al., 2021; Li et al., 2019), which is critical for lineage tracing and capturing mutant cells for downstream genomics applications.

We screened 24 *hdac1-UFlip-On* founders and identified 3 transmitting the heart BFP secondary reporter, but none of the alleles had precise 5’ or 3’ integration junctions. To recover a *hdac1-UFlip-On* allele, it may be possible to invert the entire *hdac1-UFlip-Off* cassette by Dre/*rox* mediated recombination and screen for transmission of the inverted allele in the next generation. Dre/*rox* recombination has been shown to be highly efficient in somatic tissue (Park and Leach, 2013). Consistent with this our results showed injection of Dre mRNA into *hdac1-UFlip-On* embryos led to efficient recombination and inversion of UFlip into the active orientation. Raising Dre injected embryos to identify transmitting founders may be more efficient for recovery of a *hdac1-UFlip-On* allele than re-injection to target integration of the *hdac1-UFlip-On* cassette directly.

In summary, building on our previous streamlined GeneWeld approach for CRISPR/Cas short homology directed targeted integration, here we show this method is effective for generating zebrafish floxed conditional gene alleles. We present a thorough molecular and phenotypic validation of the robustness of the UFlip floxed gene trap cassette for highly effective knockdown of gene expression and Cre/*lox* regulated conditional inactivation. UFlip alleles can be combined with any available cell or tissue specific Cre driver for spatial and temporal gene knock out, enabling the investigation of gene function in specific developmental and post embryonic stages or disease models. The UFlip vector can be adapted to introduce a variety of gene modification cassettes, such as alternative exons, synthetic exons, and complete cDNAs. In contrast to Tol2 transposon binary UAS overexpression systems, targeted integration leads to control by endogenous gene regulatory elements and should facilitate phenotypic consistency. Overall, the UFlip integration approach provides a powerful platform for generating new Cre/*lox* genetic tools and other innovations to modify gene function at endogenous loci in zebrafish.

## Materials and Methods

### Key Resources Table

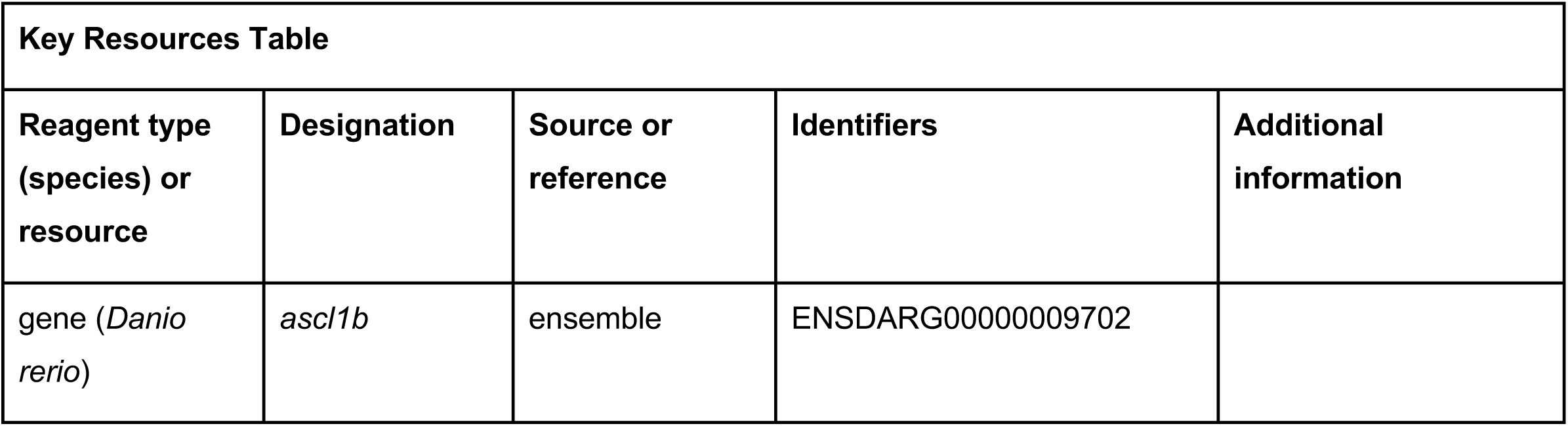
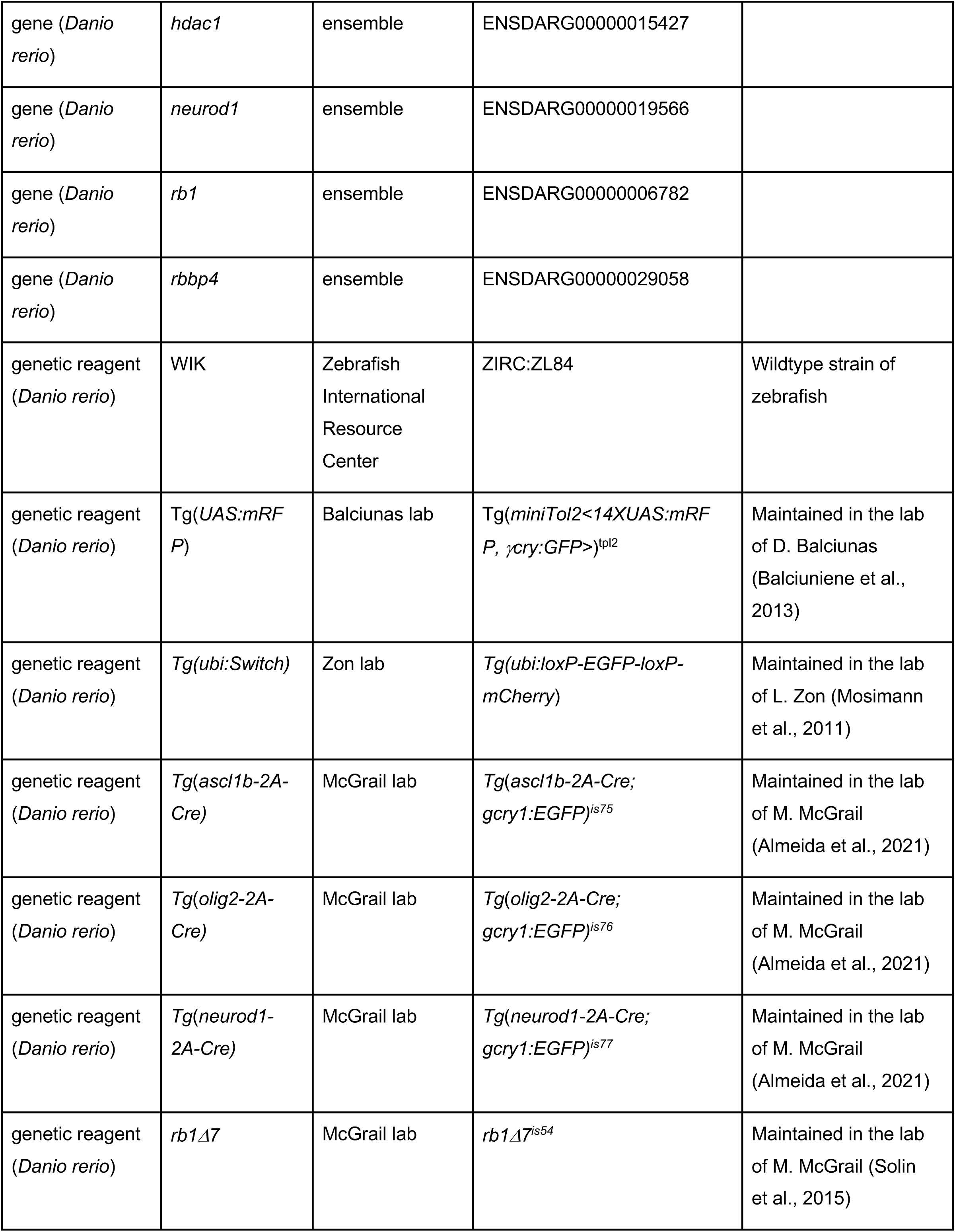
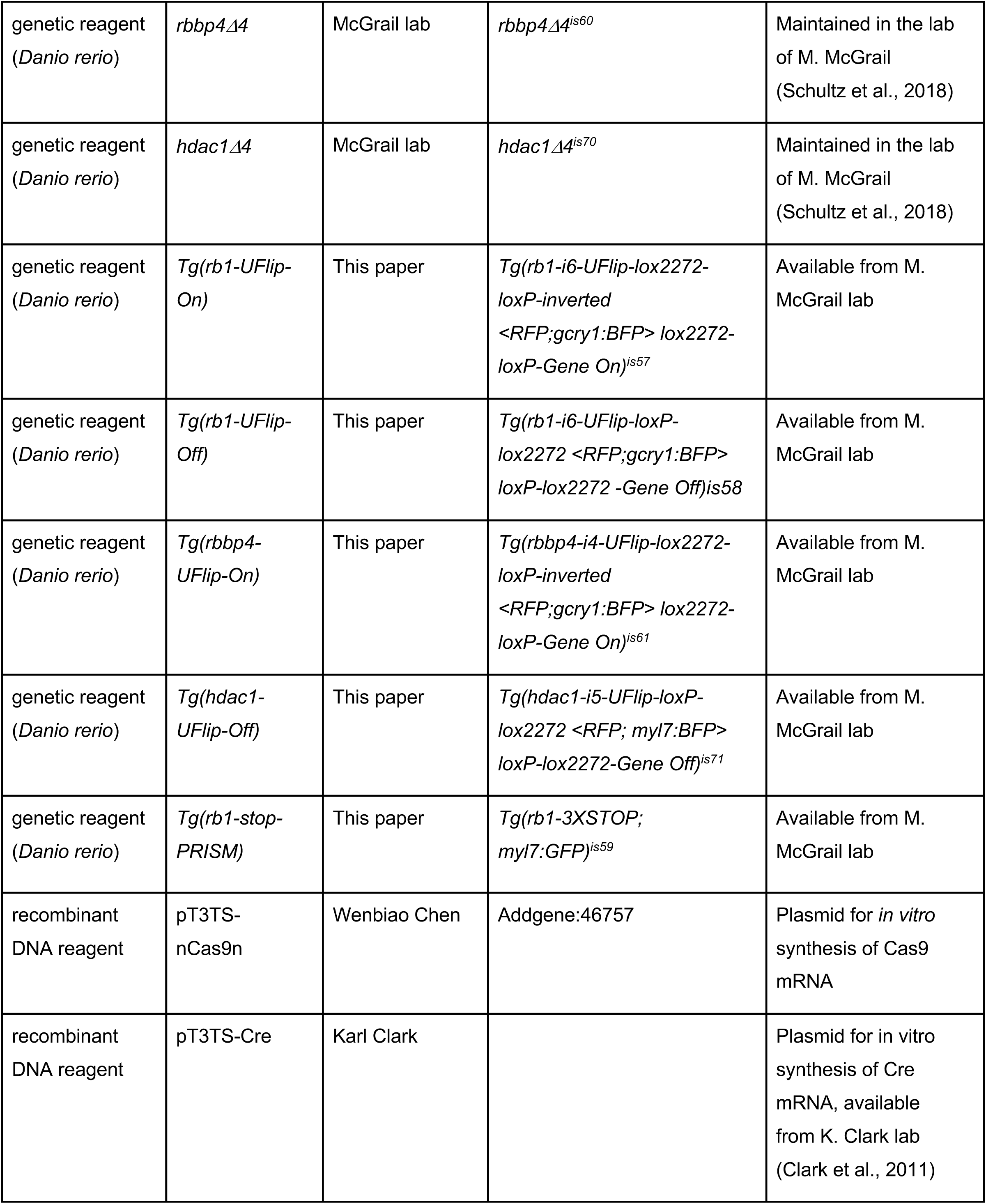
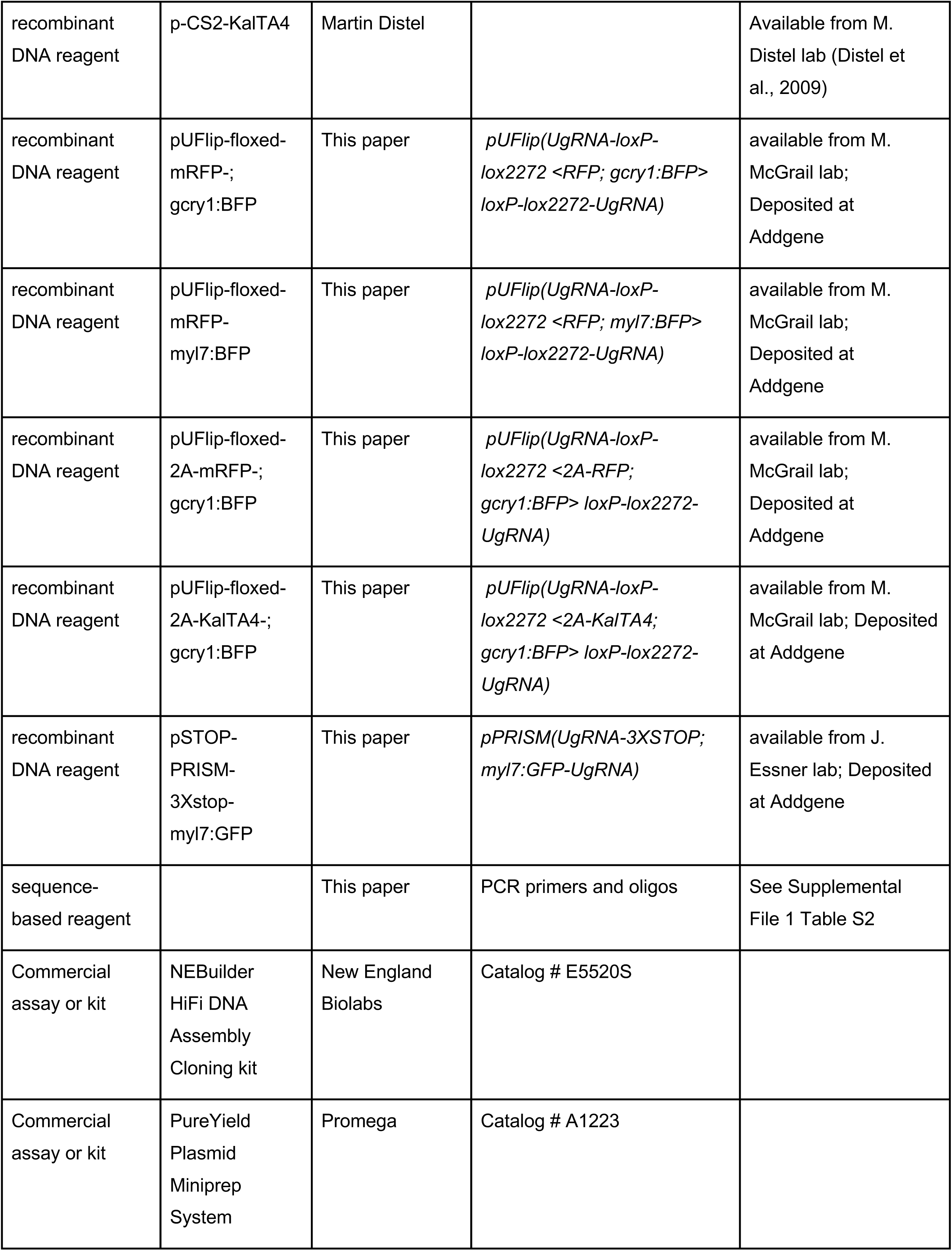
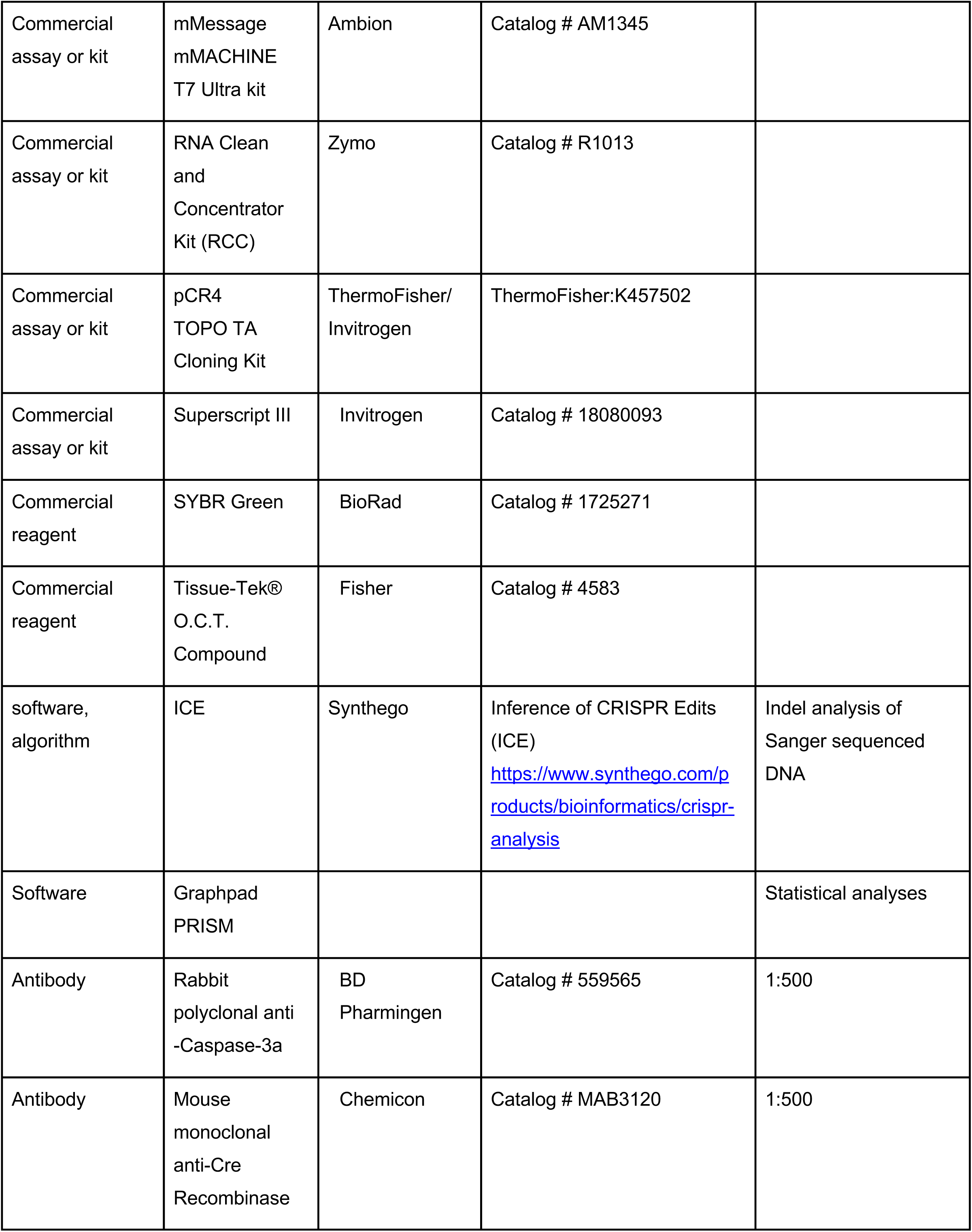
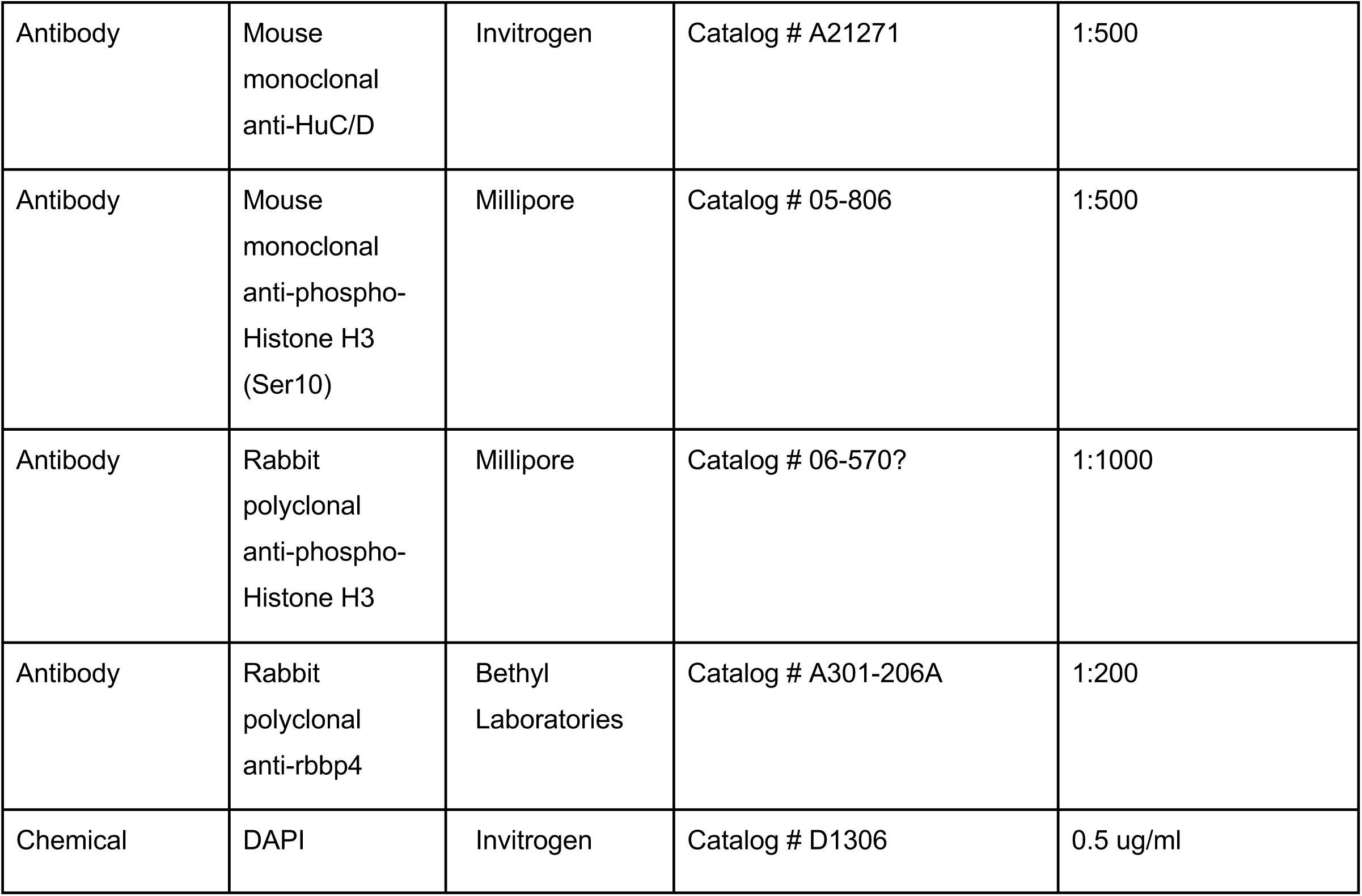

### Contact for reagent and resource sharing

Further information and requests for resources and reagents should be directed to Maura McGrail.

### Ethics Declarations and approval for animal experiments

Use of zebrafish for research in this study was performed according to the Guidelines for Ethical Conduct in the Care and Use of Animals (APA, 1986), and carried out in accordance with Iowa State University Animal Care and Use Committee IACUC-18-279 and IACUC-20-058 approved protocols. All methods involving zebrafish were in compliance with the American Veterinary Medical Association (2020), ARRIVE (Percie du Sert et al., 2020) and NIH guidelines for the humane use of animals in research.

### Zebrafish strains and maintenance

Zebrafish (*Danio rerio*) were maintained on an Aquaneering aquaculture system at 26^°^C on a 14 hour light/10 hour dark cycle. The WIK strain of wild type zebrafish was obtained from the Zebrafish International Resource Center (https://zebrafish.org/home/guide.php). Other zebrafish lines used in this study were previously described: *Tg*(*ascl1b-2A-Cre; gcry1:EGFP)*^*is75*^ and *Tg*(*neurod1-2A-Cre; gcry1:EGFP)*^*is77*^ (Almeida et al., 2021); *rb1Δ7*^*is54*^ (Solin et al., 2015); *rbbp4Δ4*^*is60*^ and *hdac1Δ4*^*is70*^ (Schultz et al., 2018); Tg(*miniTol2<14XUAS:mRFP, gcry1:GFP>*)^tl2^ (Balciuniene et al., 2013).

### Floxed UFlip vector for CRISPR-targeted integration to generate conditional alleles

The UFlip floxed mRFP gene trap vector for targeted integration (Figure 1A) was designed to be compatible with our previously published GeneWeld CRISPR/Cas9 targeted integration strategy (Wierson et al., 2020). The UFlip cassette was assembled in the pPRISM parent vector (Almeida et al., 2021) which has Universal UgRNA sequences on and BfuAI and BspQI type II restriction enzyme sites for cloning 5’ and 3’ homology arms on either side of the cassette, as previously described (Wierson et al., 2020). In the UFlip vector the homology arm cloning sites flank head to head oriented *rox* sites that sit outside alternating pairs of head to head *loxP* and *lox2272* sites for stable inversion after Cre mediated recombination (Robles-Oteiza et al., 2015; Schnutgen et al., 2003). Internal to the *rox* and *lox* sequences is the RP2 gene trap (Clark et al., 2011), consisting of a gene trap Splice Acceptor-mRFP-ocean pout (*Zoarces americanus*) antifreeze gene transcriptional termination and polyadenylation sequence (Gibbs and Schmale, 2000), followed by a tissue specific lens *gcry1:BFP* or heart *myl7:BFP* secondary reporter. To assemble UFlip an intermediate vector was created by adding EagI and ClaI restriction sites to the pPRISM backbone by PCR. 6 pairs of oligos, each containing the sequence of one of the *rox, loxP* or *lox2272* sites, with complementary overhangs were ligated into the EagI and ClaI digested backbone. All three reading frames of the RP2 gene trap plus secondary reporter was directionally cloned into AvrII and SacI restriction sites located inside the pairs of *rox*/*loxP*/*lox2272* sites. The mRFP primary reporter in the first version of the UFlip vector was subsequently replaced with the porcine teschovirus-1 polyprotein 2A peptide fused to mRFP, or KalTA4 (Distel et al., 2009) for signal amplification using the 14XUAS:RFP reporter transgenic line Tg(*miniTol2<14XUAS:mRFP, gcry1:GFP>*)^tl2^ (Balciuniene et al., 2013) (Supplemental Figures 6-8).

### Intronic sgRNA target site selection, UFlip homology arm design and targeted integration

To identify CRISPR gRNA sites in introns, intronic sequences from 4 adult female and 4 adult make WIK fish were amplified from fin clip DNA using the proofreading enzyme KOD and sequenced to identify non-repetitive sequences that were shared among adults in the population. Gene specific and vector Universal UgRNA synthetic gRNAs with 2’-O-Methyl at first and last bases, 3’ phosphorothioate bonds between first 3 and last 2 bases, were ordered from Synthego (https://orders.synthego.com/products/crisprevolution-sgrna-ez-kit-13/#/tubes?mod_code=1). gRNA efficiency was determined by co-injection of 25 pg gRNA plus 300 pg Cas9 mRNA into 1 cell stage embryos, followed by PCR amplification of the targeted intron and analysis of heteroduplex formation by gel electrophoresis. Amplicons were Sanger sequenced and the sequences analyzed for indel efficiency using Synthego’s Inference of CRISPR Edits (ICE) analysis software (https://ice.synthego.com/#/). The UFlip targeting vectors were built following the GeneWeld protocol (Welker, 2021). 24 or 48 bp 5’ and 3’ homology arms were designed to sequences flanking the genome intronic CRISPR/Cas target site. The homology arms were assembled by annealing complementary oligonucleotides with appropriate overhangs for cloning into the BfuAI and BspQI type II restriction enzymes sites that flank the UFlip cassette. To generate an active, UFlip gene off allele, the 5’ and 3’ homology arms were cloned into the UFlip BfuAI and BspQ1 type II restriction sites, respectively. To integrate the UFlip cassette in the passive, gene on orientation, the position of the cloned 5’ and 3’ homology arms was reversed, with the 3’ homology arm cloned into the BfuAI site upstream of the cassette, and the 5’ homology cloned in the BspQI site downstream of the cassette. gRNA and homology arm oligonucleotide sequences are listed in Supplementary Table S1.

For synthesis of Cas9 mRNA the expression vector pT3TS-nCas9n (Addgene #46757) (Jao et al., 2013) was linearized with Xba I (New England Biolabs, R0145S). 1 μg linearized vector was purified with the PureYield Plasmid Miniprep System (Promega, A1223) and used as template for *in vitro* synthesis of capped mRNA with the Ambion mMessage Machine T3 Transcription Kit (Thermo Fisher, AM1348). *in vitro* synthesized mRNA was purified with the RNA Clean and Concentrator Kit RCC (Zymo, R1013).

To target CRISPR/Cas driven integration of the UFlip cassette into intronic sites, a 2nl volume containing 25 pg of genomic gRNA, 25 pg of UgRNA, 10 pg of UFlip targeting vector, and 300 pg Cas9 mRNA was co-injected into 1 cell stage embryos from crosses between sequence validated adult fish. Larvae were screened at 3 dpf for expression of the lens *gcry1:BFP* or heart *myl7:BFP* secondary marker. Three to four BFP positive embryos were selected for genomic DNA isolation and confirmation of on target integration by PCR sequence analysis of 5’ and 3’ junctions. BFP positive sibling embryos were raised to adulthood. Primers, gRNA and homology arm oligonucleotide sequences are listed in Supplementary Table S1.

### Isolation of stable zebrafish *hdac1, rb1, rbbp4* UFlip and stop-PRISM integration alleles

To identify founder fish transmitting a UFlip allele, adults were outcrossed to wild type WIK and embryos were screened for expression of the UFlip secondary reporter expressing BFP in the lens (*gcry1:BFP*) or heart (*myl7:BFP*). To identify on target integration alleles, genomic DNA was extracted from individual BFP positive embryos by digestion in 50 mM NaOH at 95^°^C for 30 minutes and neutralization by addition of 1/10^th^ volume 1M Tris-HCl pH 8.0. Genomic DNA/UFlip cassette 5’ and 3’ junctions were amplified by PCR with gene specific and UFlip primers listed in Supplemental Table 2, followed by direct Sanger sequencing. BFP positive sibling embryos from a founder that was transmitting a precise UFlip integration allele were raised to adulthood. F1 adult animals were fin clipped and the genotype of individuals confirmed by PCR and sequencing. Confirmed F1 adults were outcrossed to wild type WIK, and F2 adults again confirmed for the presence of a precise UFlip integration allele. Individual confirmed F2 adults were outcrossed to WIK to establish independent transgenic lines.

The *rb1-stop-PRISM-myl7:GFP* allele was generated with a pPRISM (<u>PR</u>ecise Integration with <u>S</u>econdary <u>M</u>arker) GeneWeld targeted integration vector containing a cassette with splice acceptor followed by 3 copies of TGA, TAA, TAG, the ocean pout (*Zoarces americanus*) antifreeze gene transcriptional termination and polyadenylation sequence (Gibbs and Schmale, 2000), and a *myl7:eGFP-βactin* polyadenylation secondary reporter. Previously described 5’ and 3’ homology arms complementary to the *rb1* exon 2 CRISPR target site (Schultz et al., 2018; Wierson et al., 2020) were cloned into the BfuAI and BspQI type II restriction enzyme sites flanking the stop-PRISM-myl7:eGFP cassette. 25 pg of *rb1* exon 2 genomic gRNA, 25 pg of UgRNA, 10 pg of stop-PRISM targeting vector, and 300 pg Cas9 mRNA were coinjected into 1 cell stage embryos in a volume of 2 nl, and adult founders screened for transmission of the *rb1-stop-PRISM-myl7:GFP* allele. Precise 5’ and 3’ junctions were confirmed in heart *myl7:eGFP* expressing F2 fin clipped adults.

### Quantitative RT-PCR

Experiments to measure endogenous gene knockdown by Reverse Transcription-quantitative PCR were designed and performed according to MIQE and updated guidelines (Bustin et al., 2009; Taylor et al., 2019). Three biological replicates were performed, with each replicate representing embryos from a different mating pair of fish. At 3dpf 20 randomly selected larvae from an incross of heterozygous *hdac1-i5-UFlip-Off*/+ or *rbi-i6-UFlip-Off*/+ in-crosses were collected for RNA extraction and genotyping. Individual dissected head tissue was placed in RNA*later* (Qiagen/Thermo Fisher AM7020) or DNA/RNA shield (Zymo Research R1100-50) and individual trunk tissue was placed in 50mM NaOH for genotyping. 5 heads of each genotype were pooled and total RNA extracted using the Direct-zol RNA Microprep kit (Zymo Research, R2060) and the quality determined using a Bioanalyzer 2100 (Agilent) at the Iowa State University DNA Facility. RNA samples with a RIN>5 were normalized to the same concentration, and first strand cDNA was synthesized using SuperScript III First-Strand Synthesis SuperMix (ThermoFisher, 11752050) containing random hexamer and oligo dT primers. Primers were designed to amplify ∼200 bp amplicons with an annealing temperature of 60°C. Primer optimization and validation was performed with 3 primer concentrations (100, 200 and 400 nM) and 3 cDNA amounts (5, 25 and 215 ng) with two replicates per condition. Primer efficiency was calculated as described (Bustin et al., 2009; Taylor et al., 2019) and primer pairs with 90-100% efficiency were used for qPCR of control and test samples. The sequence of *hdac1, rb1* and reference gene *rps6kb1b* qPCR primers are listed in Supplemental Table 2. qPCR was performed on each sample in triplicate using SsoAdvanced Universal SYBR Green Supermix (Bio-Rad, 1725270) on a CFX Connect Real-Time System (Bio-Rad).

### Cre and Dre mediated inversion of UFlip alleles

For synthesis of Cre and Dre mRNAs, the expression vectors pT3TS-Cre (Clark et al., 2011) was linearized with SalI (New England Biolabs, R0138S) and pT3TS-Dre was linearized with BamHI (New England Biolabs, R0136S). 1 μg linearized vector was purified with the PureYield Plasmid Miniprep System (Promega, A1223) and used as template for *in vitro* synthesis of capped mRNA with the Ambion mMessage Machine T3 Transcription Kit (Thermo Fisher, AM1348). *in vitro* synthesized mRNA was purified with the RNA Clean and Concentrator Kit RCC (Zymo, R1013). 12.5 pg Cre or 15 pg Dre mRNA was injected into 1 cell stage embryos to promote recombination mediated inversion of the UFlip cassette at *lox* or *rox* sites. UFlip cassette inversion in Cre or Dre injected 3 dpf larvae was confirmed by digestion of individual larvae in 50 mM NaOH at 95^°^C for 30 minutes and neutralization by addition of 1/10^th^ volume 1M Tris-HCl pH 8.0. Genomic DNA/UFlip cassette 5’ and 3’ junctions were amplified by PCR with gene specific and UFlip primers listed in Supplemental Table 2, followed by direct Sanger sequencing.

### Zebrafish tissue embedding, sectioning, immunolocalization and imaging

Zebrafish embryo and larvae fixation, embedding, sectioning and immunolabeling was as described previously (Schultz et al., 2018). Zebrafish embryos and larvae were anesthetized in 160ug/ml Ethyl 3-aminobenzoate methanesulfonate (Tricaine, MS-222) C9H11NO2·CH4SO3 (Sigma-Aldrich, 886-86-2) in E3 embryo media (Westerfield, 1995) and head and trunk dissected. Trunk tissue was placed in 20µl 50mM NaOH for genotyping. Heads were fixed in 4% paraformaldehyde overnight at 4°C, incubated in 30% sucrose overnight at 4°C, then processed and embedded in Tissue-Tek OCT (Fisher, 4583). Tissues were sectioned at 14-16 µm on a Microm HM 550 cryostat. Antibodies used for labeling: rabbit polyclonal anti-phospho-Histone H3 PH3 1:1000 (Cell Signaling Technology; 9701); mouse monoclonal anti-phospho-Histone H3 (Ser10), clone 3H10 1:500 (Millipore 05-806); mouse monoclonal anti-HuC/D 1:500 (Invitrogen A-21271); mouse monoclonal Anti-Cre recombinase 1:250 (Millipore-Sigma MAB3120); rabbit polyclonal anti-Caspase-3a 1:500 (BD Biosciences 559565); Alexa-594 (Invitrogen A-11005) and Alexa-488 (Invitrogen A-11008) conjugated secondary antibodies 1:500. Tissues were counterstained with 5 µg/ml DAPI, mounted in Fluoro-Gel II containing DAPI (Electron Microscopy Sciences 17985-50) and imaged on a Zeiss LSM700 or LSM800 laser scanning confocal microscope.

For live imaging, embryos and larvae were treated with 0.003% 1-phenyl 2-thiourea (Sigma, P7629) to inhibit pigment synthesis, anesthetized in Tricaine in embryo media (Westerfield, 1995) and mounted on slides in 1.2% low-melt agarose/embryo media/Tricaine. Fluorescence and bright field imaging were performed on a Zeiss SteREO Discovery V12 microscope equipped with an X-Cite 120W Metal Halide lamp (Excilitas Technologies, X-Cite 120Q). Images were captured with a Cannon Rebel T3 camera using EOS Utility software (Cannon). Bright field and fluorescence images were merged in Photoshop (Adobe).

### Quantification and statistical analyses

Quantification of proliferation and apoptosis was performed on 3 sections of immunolabeled head tissue for each individual, from 3 biological replicates of zebrafish embryos or larvae. Prism (GraphPad) software was used for two-tailed unpaired Student’s t-test with mean ± s.e.m. statistical analyses and production of bar graphs.

## Data Availability

All DNA constructs reported in this study have been deposited at Addgene in the Jeffrey Essner lab list. DNA constructs and transgenic zebrafish lines are available on request to M. McGrail.

## Acknowledgements

The authors thank Dr. Darius Balciunas for the transgenic line Tg(*miniTol2<14XUAS:mRFP, gcry1:GFP>*)^tl2^ and Dr. Raquel Espin (Iowa State University) for the KalTA4 cDNA. This work was supported by NIH grants R24OD020166 (MM, JJE, DLD, IF, KJC, SCE) and by a graduate scholarship from the CNPq Brazilian National Council for Scientific and Technological Development (MPA).

## Author Contributions

MPA, JMW, JJE and MM conceived the study; MPA and MM wrote the manuscript with input from JJE, SK, FL, JMW, WAW, SCE and KJC; MPA, SK, FL, JMW, WAW, ZM and LES-R designed and performed the experiments; JMW, ZM, KJC, JJE and MM designed and built the UFlip vectors.

## Figure Legends

**Supplemental Figure 1.**
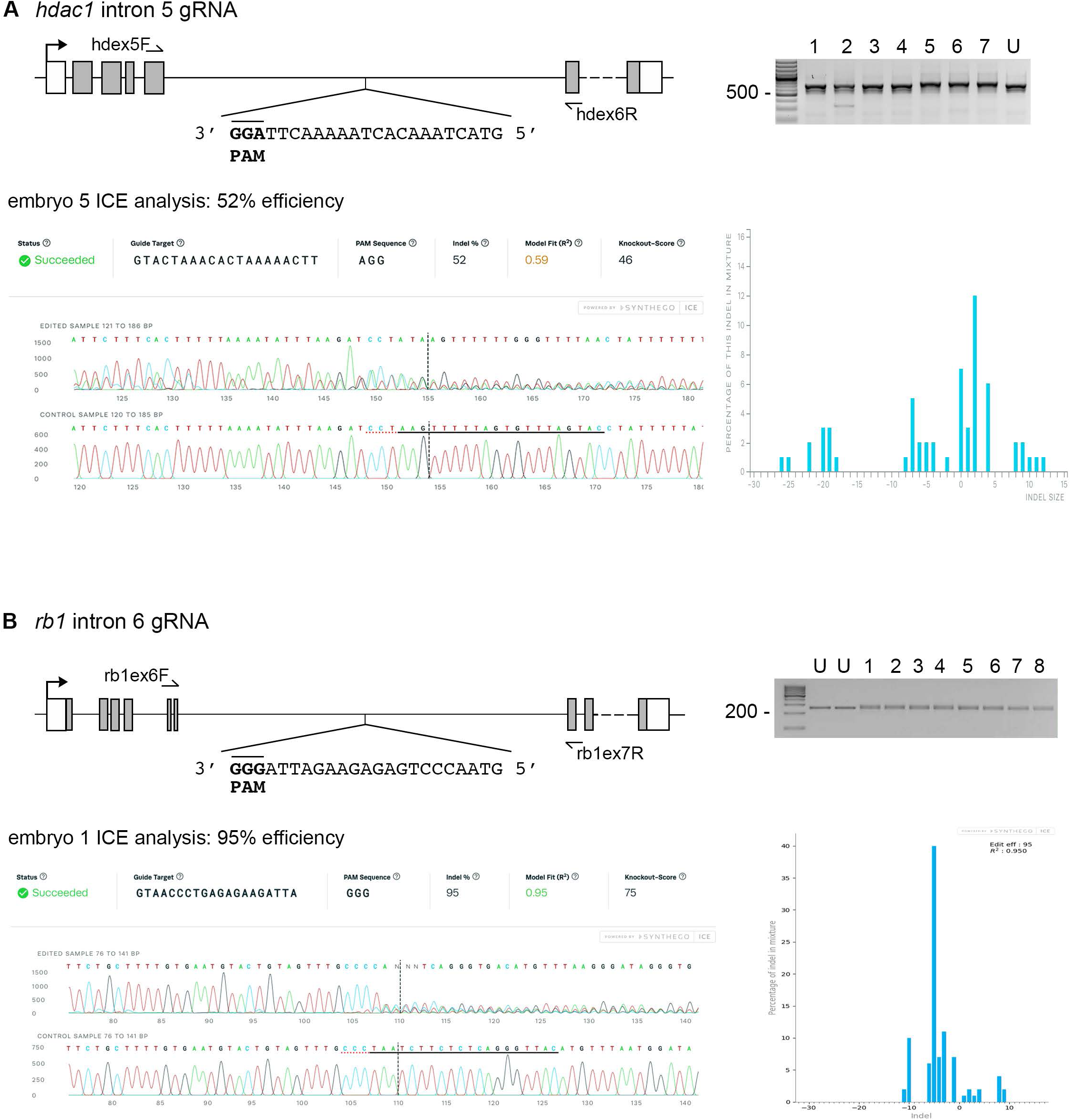
ICE analysis to test *hdac1* and *rb1* intronic gRNA efficiency. (**A**) *hdac1* gene model with sequence of the intron 5 reverse strand gRNA. Gel image of PCR amplicons surrounding the target site from 7 embryos injected with Cas9 and the gRNA (1-7), and 1 uninjected embryo (U). Amplicons from embryo #5 and the uninjected embryo were sequenced and analyzed with Synthego’s ICE software, and indicate 52% indel efficiency at the target site. (**B**) *rb1* gene model with sequence of the intron 6 reverse strand gRNA. Gel image of PCR amplicons surrounding the target site from 8 embryos injected with Cas9 and the gRNA (1-8), and 2 uninjected embryos (U). Amplicons from embryo #1 and an uninjected embryo were sequenced and analyzed with Synthego’s ICE software, and indicate 95% indel efficiency at the target site. Plots show the range and percentage of indels present in the sequences. PAM sequences shown in bold and underlined.

**Supplemental Figure 2.**
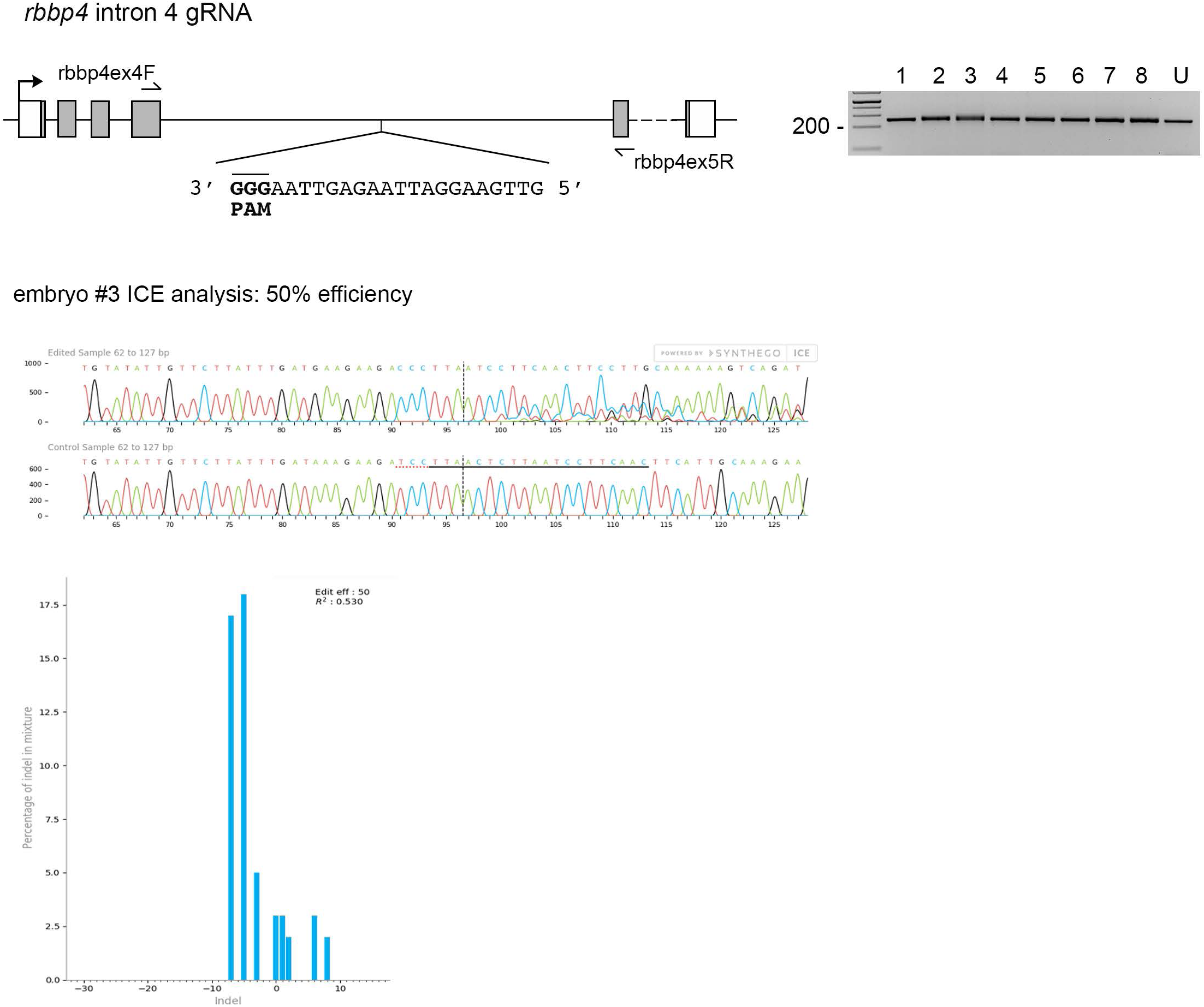
ICE analysis to test *rbbp4* intronic gRNA efficiency. *rbbp4* gene model with sequence of the intron 4 reverse strand gRNA. Gel image of PCR amplicons surrounding the target site from 8 Cas9 plus gRNA injected and 1 uninjected (U) embryo. Amplicons from embryo #3 and the uninjected embryo were sequenced and analyzed with Synthego’s ICE software, and indicate 50% indel efficiency at the target site. Plot shows the range and percentage of indels present in the sequences. PAM sequence shown in bold and underlined.

**Supplemental Figure 3.**
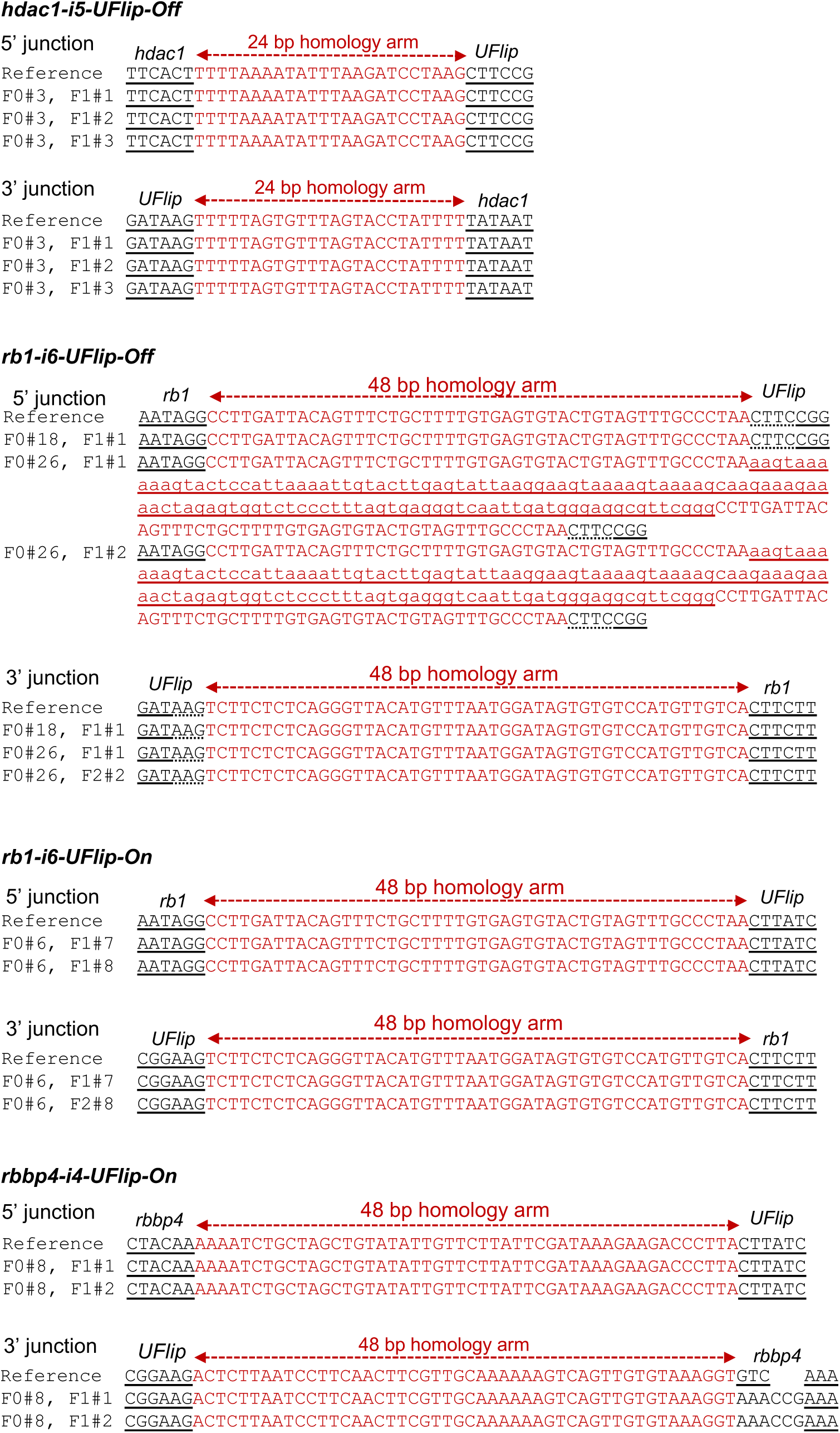
*hdac1-UFlip-Off, rb1-UFlip-Off, rb1-UFlip-On* and *rbbp4-UFlip-On* integration allele genomic DNA/UFlip junction analysis. 5’ and 3’ genomic-UFlip integration junctions were PCR amplified from F1 transgenic zebrafish fin clip genomic DNA. The PCR products were sequenced and aligned to the reference sequence expected for a precise integration at the genomic target site. Capitalized red nucleotides represent 24 or 48 bp homology arms. *rb1-UFlip-Off* founder #26 transmitted an allele with a 5 junction that included a duplication of the 5’ Homology Arm sequence that flanked a 115 bp segment of the vector backbone (lowercase underlined nucleotides). The *rbbp4-UFlip-On* 3’ junction contained a 3 bp deletion plus 6 bp insertion between the *rbbp4* 48 bp Homology Arm and flanking genomic DNA sequences.

**Supplemental Figure 4.**
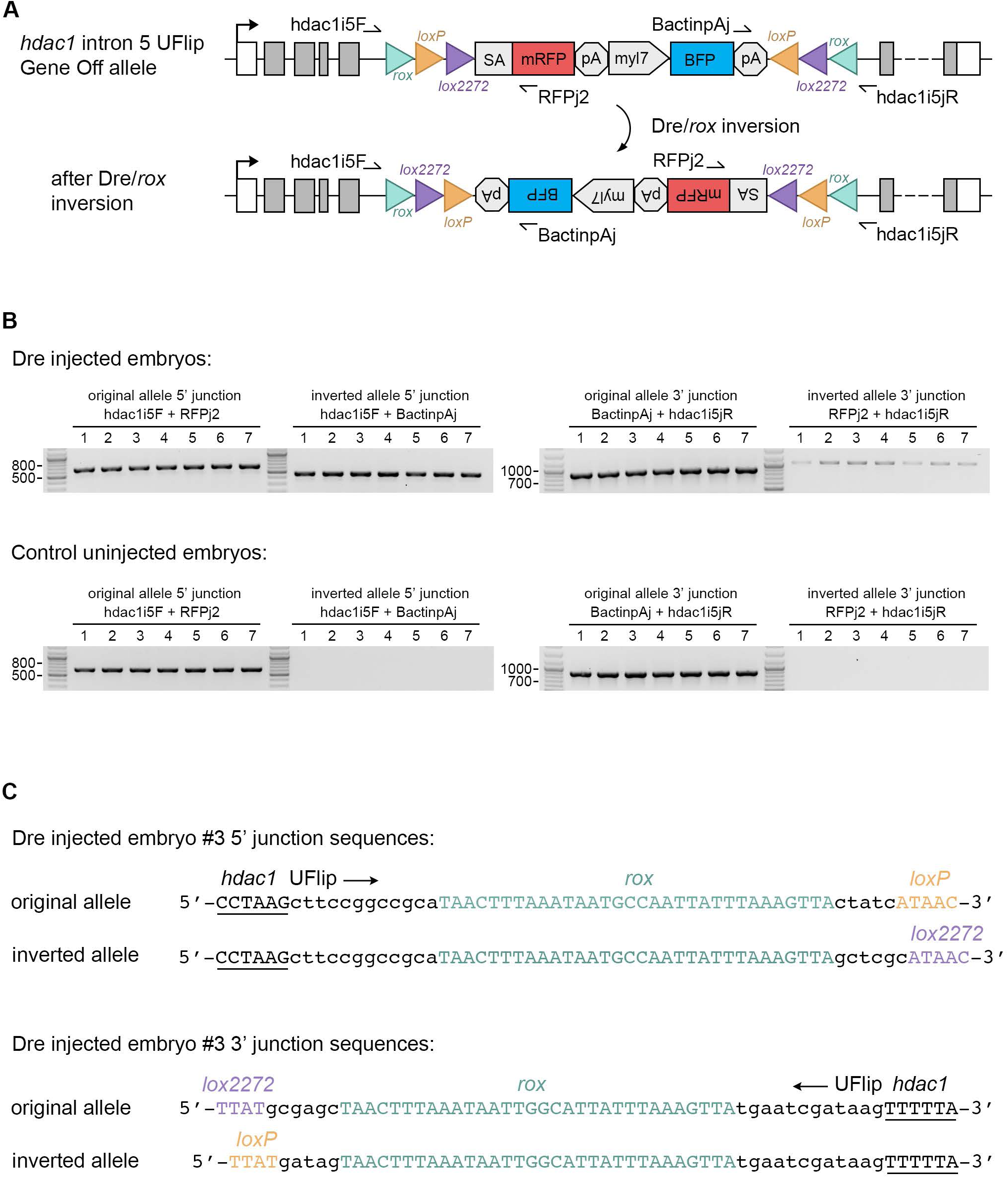
Dre/*rox*-mediated inversion of the *hdac1-UFlip-Off* cassette after Dre mRNA injection into embryos. (**A**) Diagram of *hdac1-UFlip-Off* allele before and after Dre/*rox* inversion. Primer pairs for 5’ and 3’ junctions before and after inversion are shown. (**B**) 7 *hdac1-UFlip-Off/+* Dre injected embryos from an *hdac1-UFlip-Off/+* outcross to wild type WIK were analyzed by PCR with primers to the original and inverted 5’ and 3’ junctions. Each of the injected embryos resulted in amplification of the original and inverted cassette 5’ and 3’ junction bands. Uninjected controls only showed amplification of the original gene off allele 5’ and 3’ junctions. (**C**) Sequence analysis of original and inverted junction PCR products confirmed Dre injection led to cassette inversion at the *rox* sites to the gene on orientation.

**Supplemental Figure 5.**
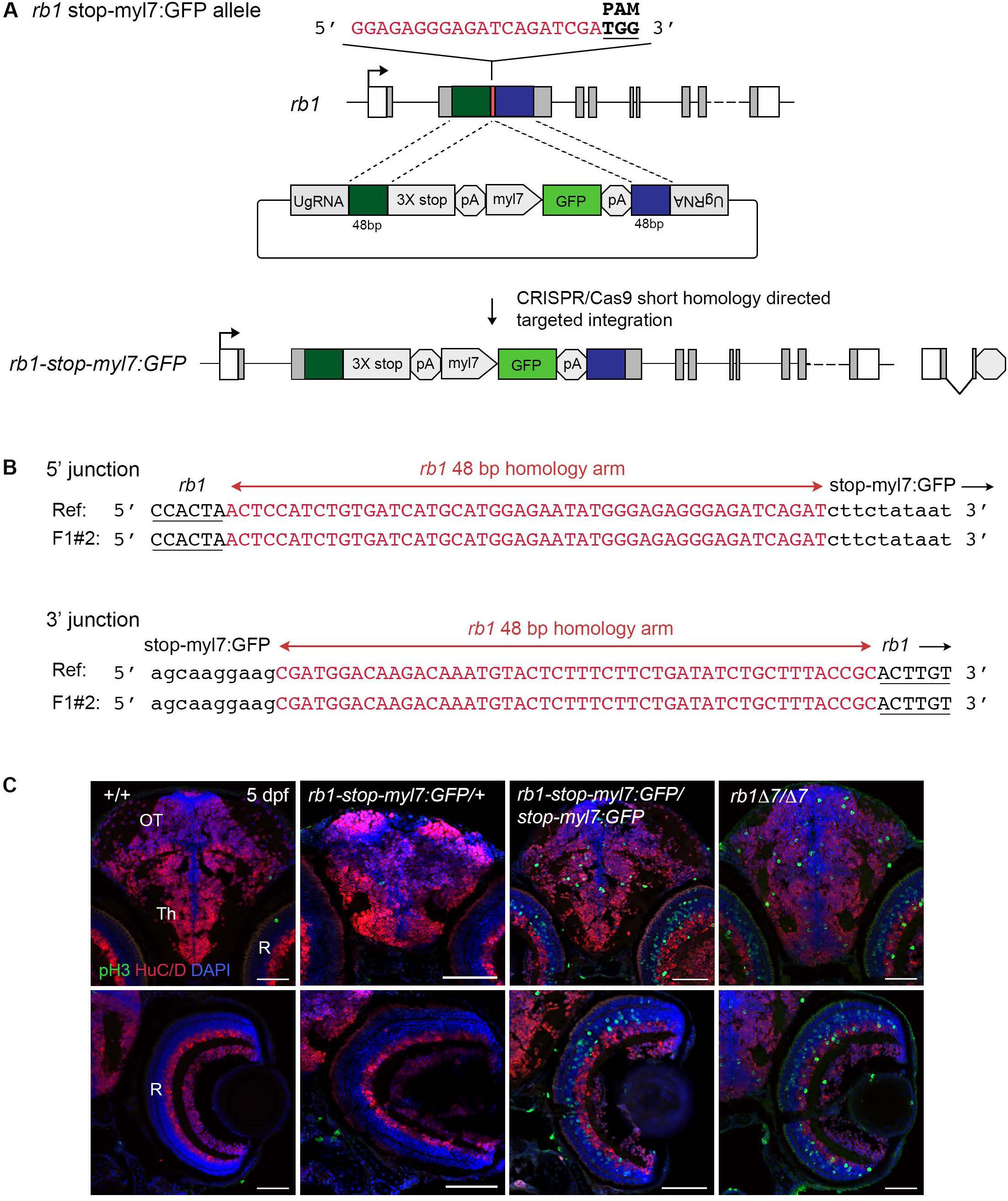
The *rb1-stop* integration allele recapitulates the *rb1D7* indel loss of function phenotype. (**A**) Diagram of *rb1* gene model with exon 2 gRNA, the stop-PRISM-myl7:GFP targeting vector, and the resulting *rb1-stop-myl7:GFP* allele after GeneWeld CRISPR/Cas9 targeted integration. (**B**) F1 adult *rb1-stop* 5’ and 3’ junction analysis shows precise integration of the stop-PRISM cassette at the exon 2 target site. (**C**) Immunolocalization control +/+, *rb1-stop/+, rb1-stop/rb1-stop/*, and *rb1D7*/*rb1D7* 5 dpf sectioned head tissue shows proliferating pH3 positive cells throughout the midbrain optic tectum and thalamic region (top row) and retina (bottom row) in both *rb1-stop-myl7:GFP* and *rb1D7* homozygotes. Th, thalamic region; OT, optic tectum; R, retina. Scale bars: 50 μm (**C**).

**Supplemental Figure 6.**
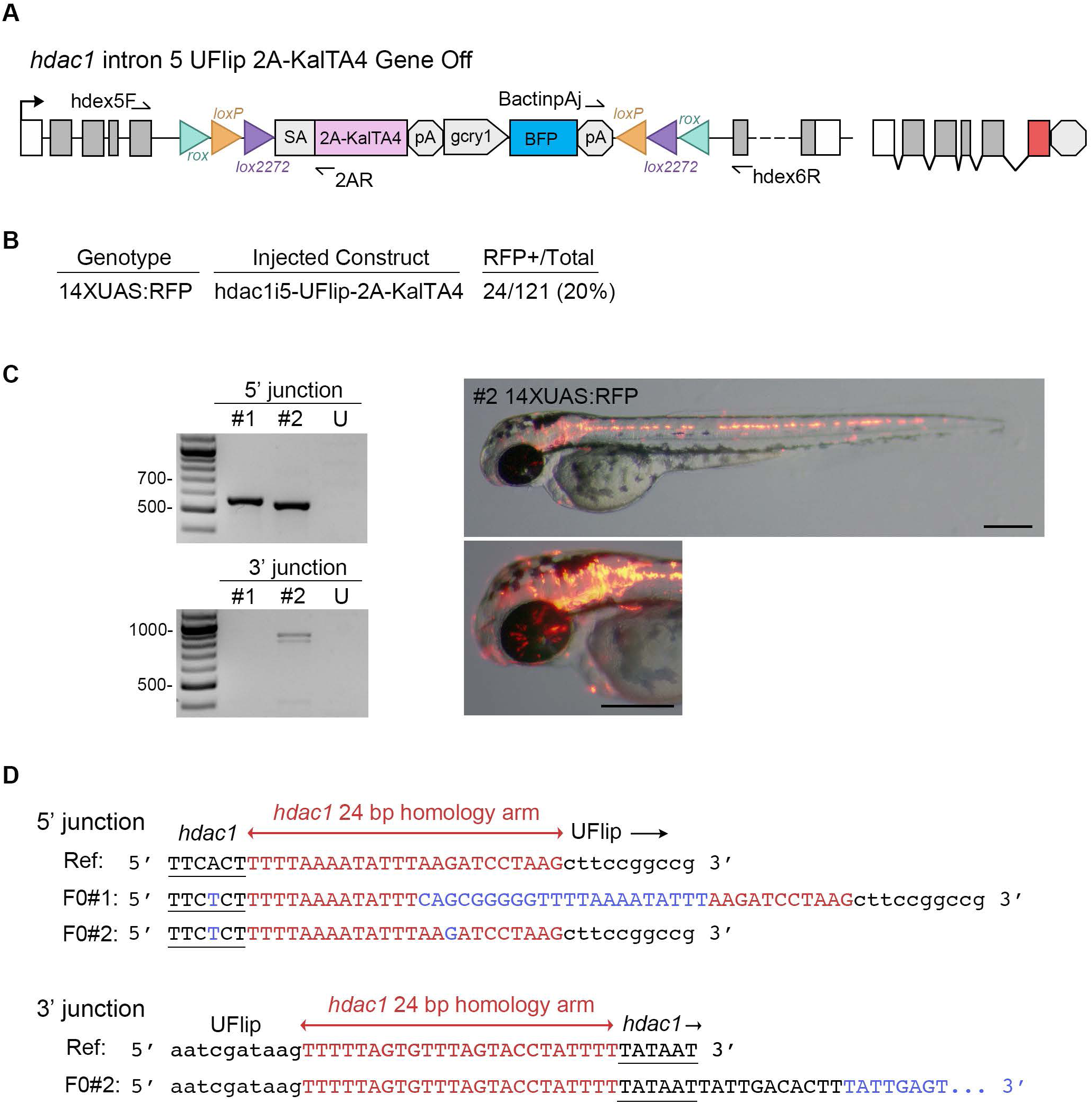
Targeted integration of the UFlip-2A-KalTA4 cassette into *hdac1* intron 5 leads to robust expression of the primary reporter. (**A**) Diagram of UFlip-2A-KalTA4-Off integration into *hdac1*. (**B**) 20% of 14XUAS:mRFP injected embryos show mRFP expression. (**C**) Junction fragment analysis from 2 injected mRFP positive embryos. Embryo #2 showed 5’ and 3’ junctions, and widespread expression throughout the nervous system. (**D**) Sequence analysis of junction fragments from targeted embryos reveals the 5’ junction in #1 has a 22 base pair insertion within the 24 bp 5’ homology arm. The 3’ junction in individual #2 has 17 base pairs matching the reference genome, followed by a 67 base pair insertion. The 67 base pair insertion most likely represents a polymorphism in the WIK population. Scale bar: 200 μm (**C**).

**Supplemental Figure 7.**
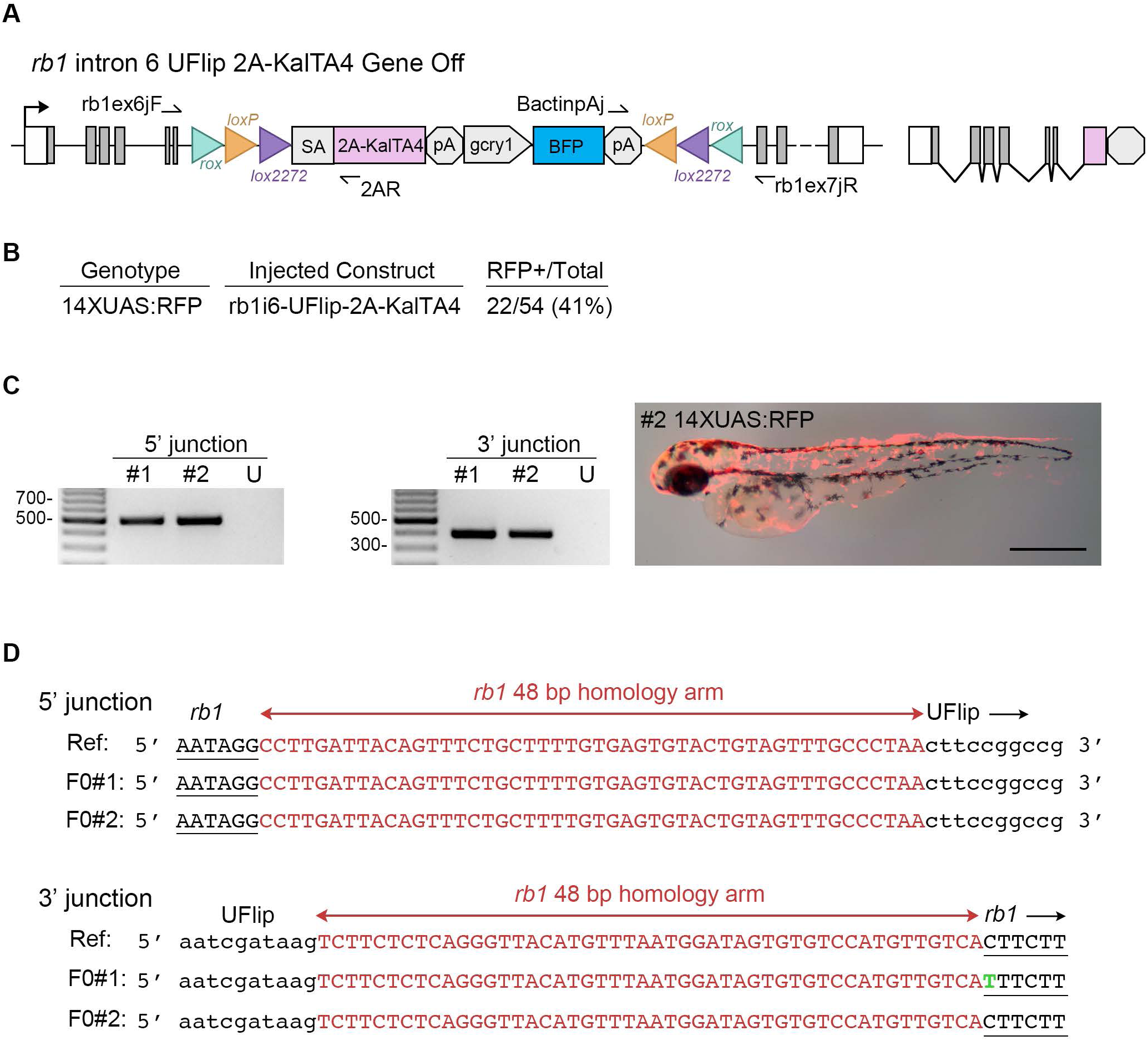
Targeted integration of the UFlip-2A-KalTA4 cassette into *rb1* intron 6 leads to robust expression of the primary reporter. (**A**) Diagram of UFlip-2A-KalTA4-Off integration into *rb1*. (**B**) 41% of 14XUAS:mRFP injected embryos show mRFP expression. (**C**) Junction fragment analysis from 2 injected mRFP positive embryos. Both embryos showed 5’ and 3’ junctions. Image shows widespread mRFP expression throughout embryo #2. (**D**) Sequence analysis of junction fragments from targeted embryos reveals 5’ and 3’ precise on target integration. Scale bar: 200 μm (**C**).

**Supplemental Figure 8.**
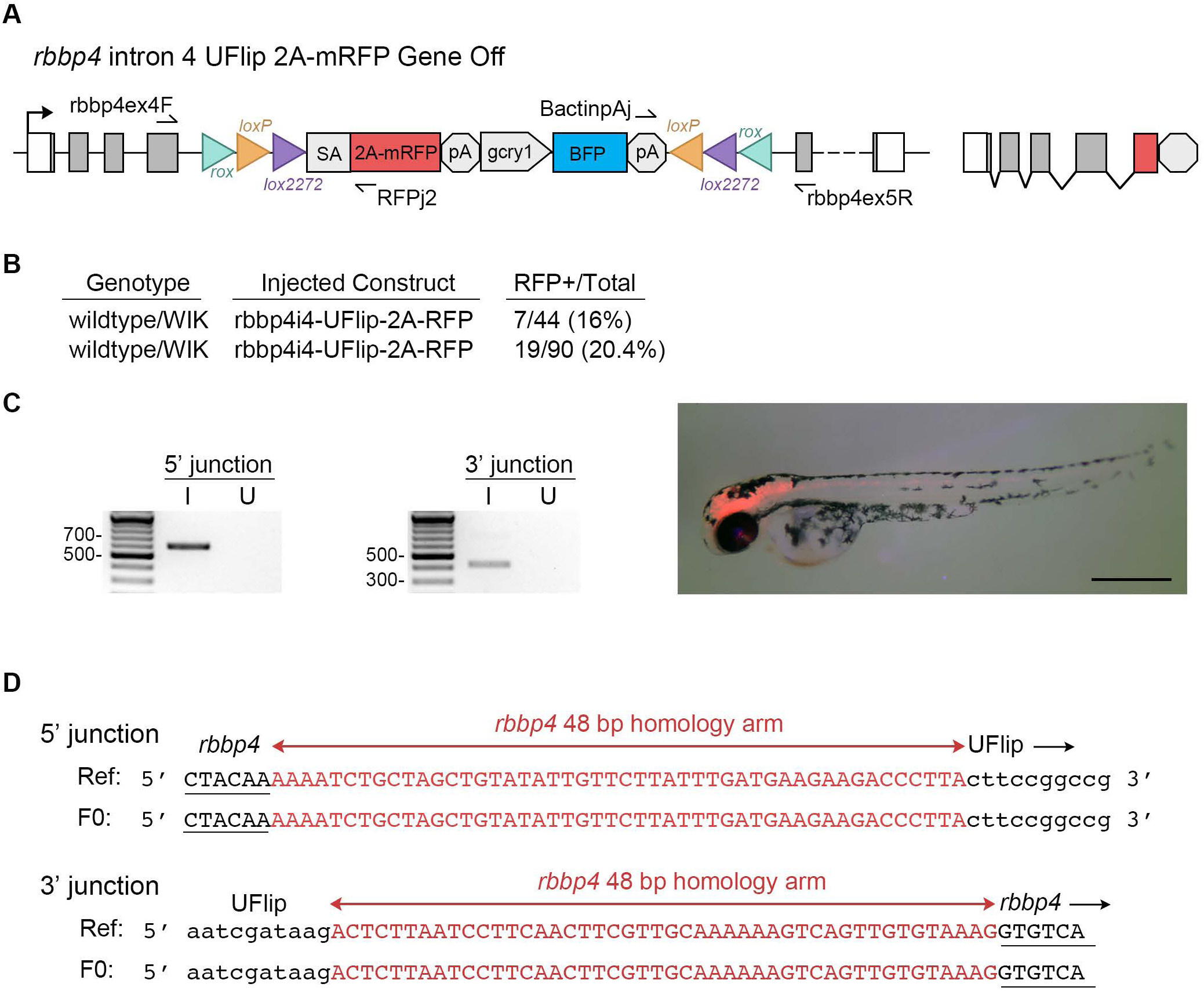
Targeted integration of the UFlip-2A-mRFP cassette into *rbbp4* intron 4 leads to robust expression of the primary reporter. (**A**) Diagram of UFlip-2A-mRFP-Off integration into *rbbp4*. (**B**) 16% and 20.4% of 14XUAS:mRFP injected embryos from two separate injections show mRFP expression. (**C**) Junction fragment analysis from 1 injected mRFP positive embryo showed 5’ and 3’ junctions. Image shows mRFP expression throughout the central nervous system. (**D**) Sequence analysis of junction fragments from the mRFP positive embryo in **C** reveals 5’ and 3’ precise on target integration. Scale bar: 200 μm (**C**).

**Supplemental Figure 9.**
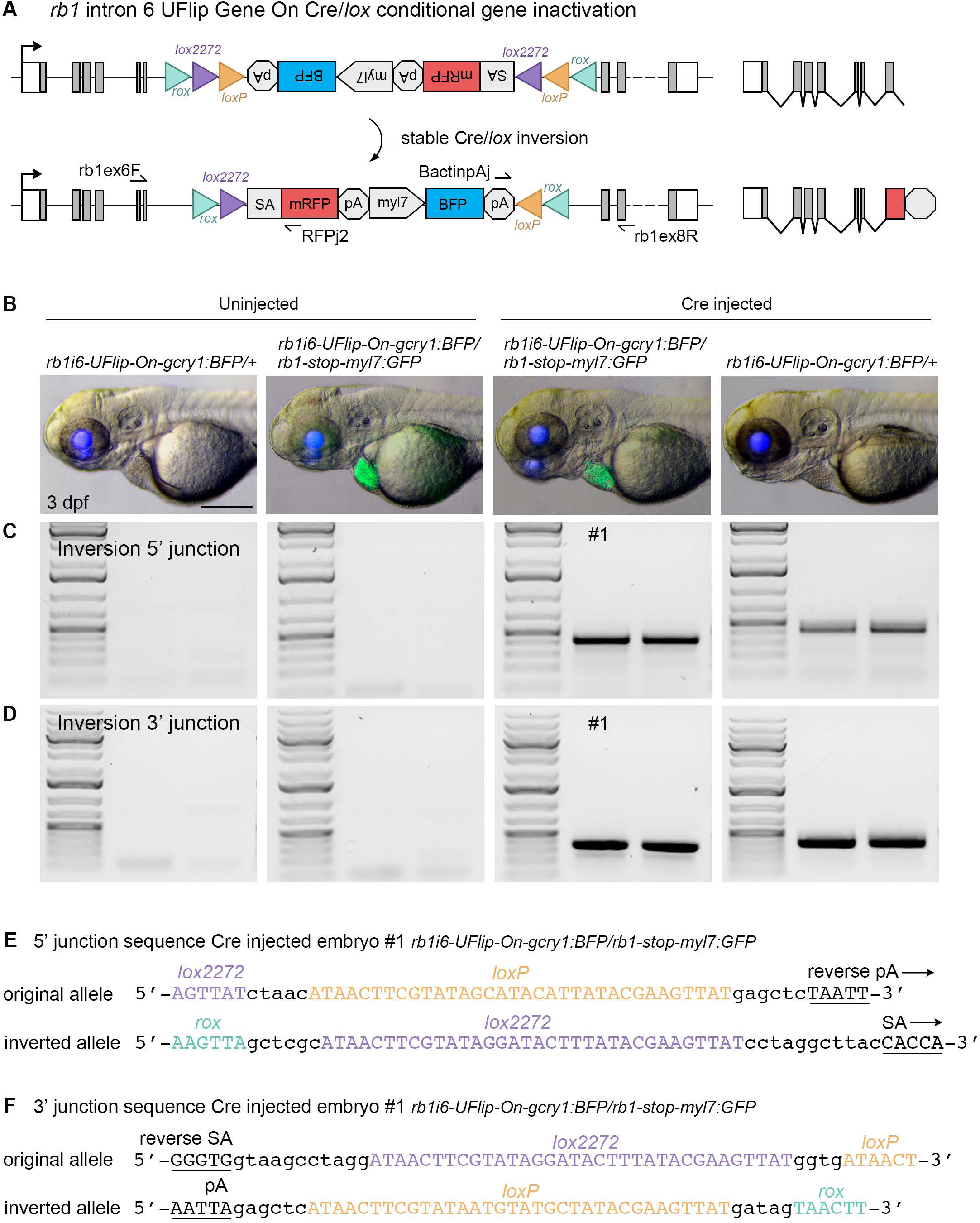
Validation of Cre/*lox*-mediated *rb1-UFlip-On* inversion after Cre mRNA injection into embryos. (**A**) Diagram of *rb1-UFlip-On* allele before and after stable Cre/*lox* inversion. Location of primers used for PCR amplification of genomic DNA/UFlip junction and inversion are indicated. (**B**) Gross phenotype of uninjected and Cre mRNA injected *rb1-UFlip-On/+* and *rb1-UFlip-On/rb1-stop* 3 dpf larvae. (**C, D**) PCR analysis of uninjected and Cre mRNA injected *rb1-UFlip-On/+* and *rb1-UFlip-On/rb1-stop* embryos to test for 5’ junction (**C**) and 3’ junction (**D**) after Cre/*lox* inversion. (**E, F**) Sequence analysis of PCR amplicons from Cre injected *rb1-UFlip-On/rb1-stop* embryo #1 confirms Cre/*lox* mediated inversion of the UFlip cassette. Scale bar: 200 μm (**B**).

**Supplemental Figure 10.**
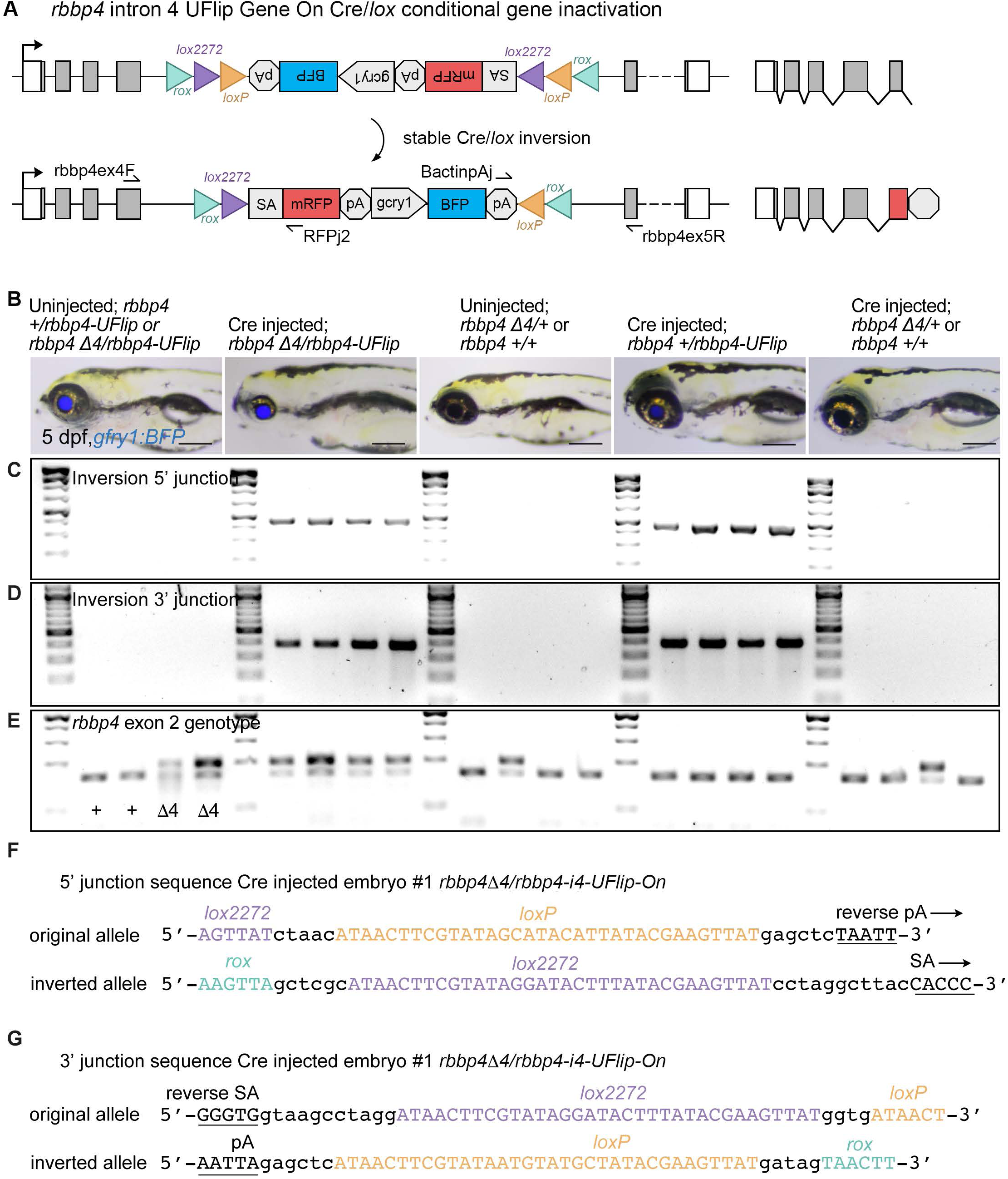
Validation of Cre/*lox*-mediated *rbbp4-UFlip-On* inversion after Cre mRNA injection into embryos. (**A**) Diagram of *rbbp4-UFlip-On* allele before and after stable Cre/*lox* inversion. Location of primers used for PCR amplification of genomic DNA/UFlip junction and inversion are indicated. (**B**) Gross phenotype of uninjected and Cre mRNA injected 5 dpf larvae from adult *rbbp4D4*/+ crossed to *rbbp4-UFlip-On/+*. Larvae were sorted based on expression of the *rbbp4-UFlip* secondary marker lens BFP and the *rbbp4* loss of function microcephaly and microphthalmia phenotype. (**C, D**) PCR analysis of 4 uninjected and Cre mRNA injected larvae from each class to test for 5’ junction (**C**) and 3’ junction (**D**) after Cre/*lox* inversion. (**E**) PCR genotyping confirmation of *rbbp4* wild type and *rbbp4D4* alleles. (**F, G**) Sequence analysis of PCR amplicons from Cre injected *rbbp4D4*/*rbbp4-UFlip-On* embryo #1 confirms Cre/*lox* mediated inversion of the UFlip cassette. Scale bar: 200 μm (**B**).

---

**Supplementary Table 1.**
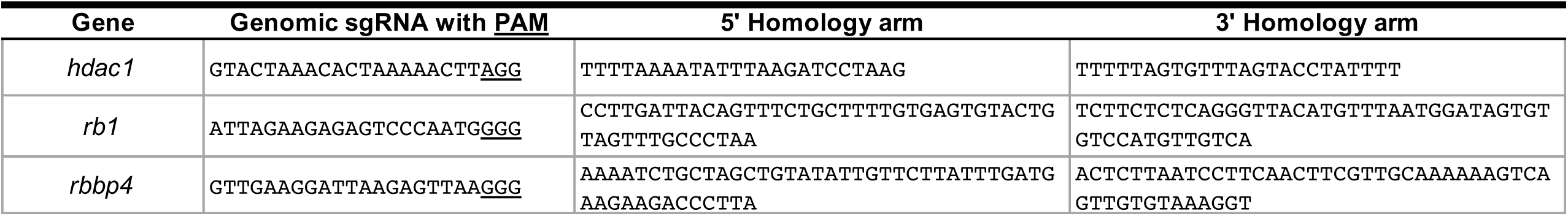
Genome intronic CRISPR gRNA sites and UFlip targeting vector homology arm sequences.

**Supplementary Table 2.**
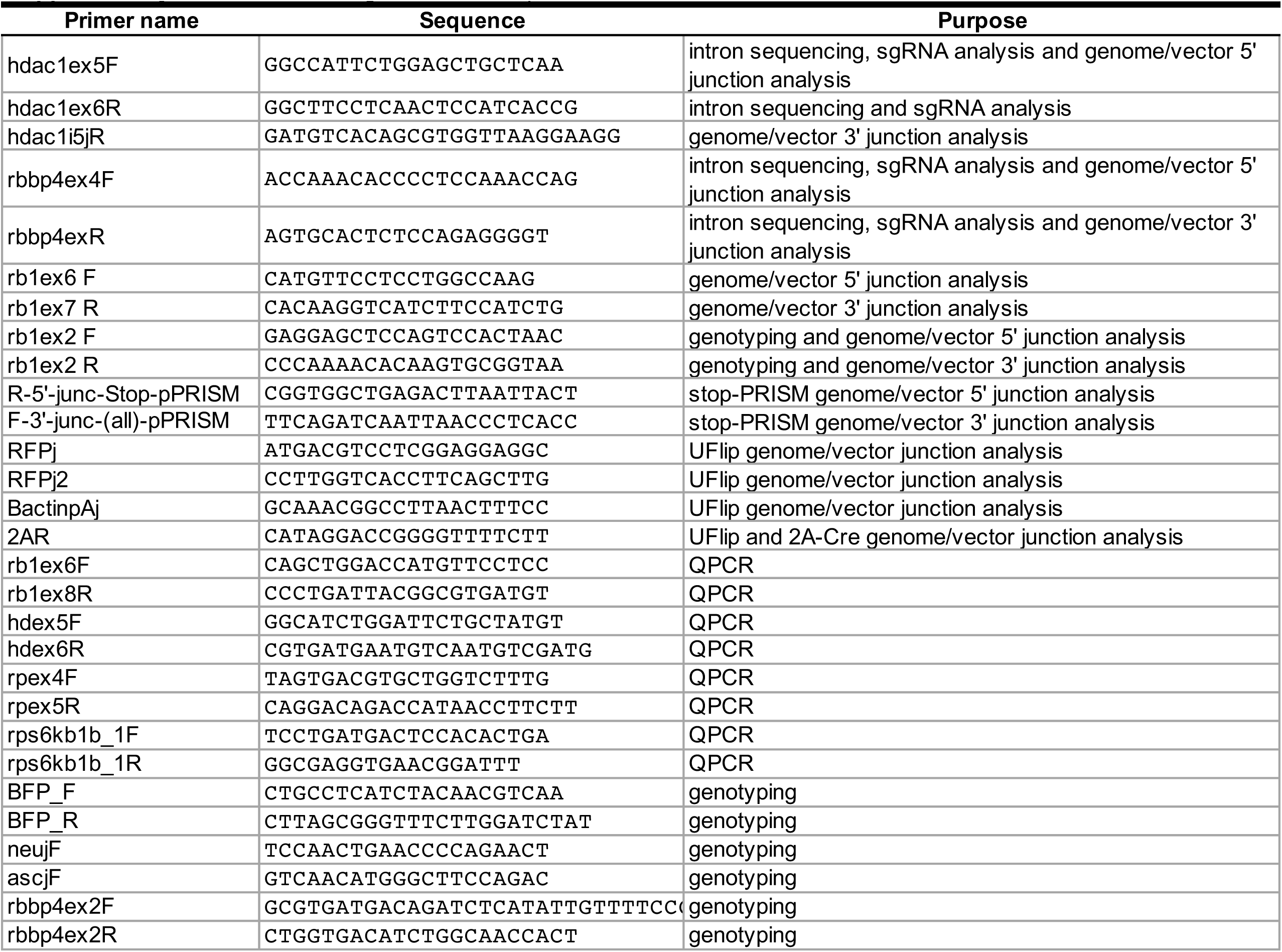
Primer Oligonucleotide sequences.

